# Desmosome mutations impact the tumor microenvironment to promote melanoma proliferation

**DOI:** 10.1101/2023.09.19.558457

**Authors:** Maayan Baron, Mohita Tagore, Patrick Wall, Fan Zheng, Dalia Barkley, Itai Yanai, Jing Yang, Maija Kiuru, Richard M. White, Trey Ideker

## Abstract

Desmosomes are transmembrane protein complexes that contribute to cell-cell adhesion in epithelia and other tissues. Here, we report the discovery of frequent genetic alterations in the desmosome in human cancers, with the strongest signal seen in cutaneous melanoma where desmosomes are mutated in >70% of cases. In primary but not metastatic melanoma biopsies, the burden of coding mutations in desmosome genes associates with a strong reduction in desmosome gene expression. Analysis by spatial transcriptomics and protein immunofluorescence suggests that these expression decreases occur in keratinocytes in the microenvironment rather than in primary melanoma cells. In further support of a microenvironmental origin, we find that desmosome gene knockdown in keratinocytes yields markedly increased proliferation of adjacent melanoma cells in keratinocyte/melanoma co-cultures. Similar increases in melanoma proliferation are observed in media preconditioned by desmosome-deficient keratinocytes. Thus, gradual accumulation of desmosome mutations in neighboring cells may prime melanoma cells for neoplastic transformation.

## Introduction

Transformation of melanocytes to skin cutaneous melanoma (SKCM) includes interaction with numerous cells in the tumor microenvironment which can play important roles in progression^1–3^. One important component of this microenvironment is the keratinocyte, which forms the major cellular component of the epidermis, the top layer of skin. Nascent melanoma cells detach from neighboring keratinocytes, regulating melanocyte homeostasis through paracrine signaling and cell-cell adhesion^4–10^ and impacting tumor proliferation and invasion.

A primary adhesion structure that facilitates the physical interaction between cells in the epithelium is the desmosome, an intercellular junction conserved across vertebrates^11^. Desmosomes have a transmembrane core composed of cadherin proteins (desmogleins and desmocollins) which is linked via plakoglobin and plakophilin proteins to keratin filaments in the cytoplasm^12^ (Fig. 1a). Alterations to desmosome components can impair tissue strength and have been documented to have both positive and negative effects on cell proliferation and differentiation^13^. For example, increases in expression of desmosome cadherins can stimulate release of plakoglobin followed by increased Wnt/β-catenin signaling, and overexpression of plakophilin 3 (PKP3) has been associated with poor patient survival and advanced disease stage in non-small cell lung carcinomas^14^. In contrast, upregulation of plakoglobin has been found to suppress cell proliferation in bladder cancer cells *in vitro*^15,16^.

**Fig. 1:**
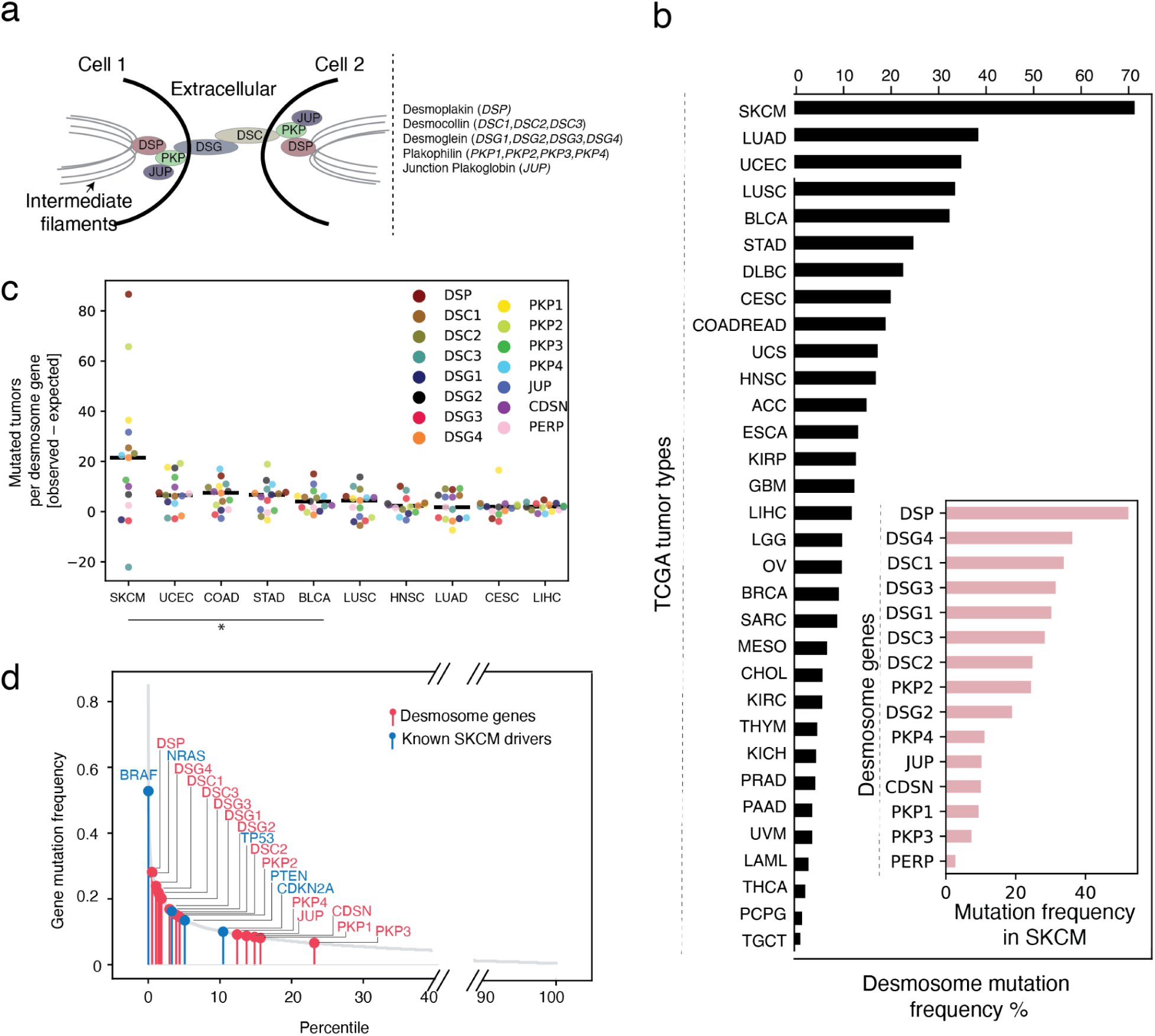
The desmosome system and its patterns of mutation. (a) Illustration of the desmosome complex, which is composed of proteins of three different families: cadherins (*DSG*s and *DSC*s), armadillo proteins (*PKP*s and *JUP*) and the plakin protein desmoplakin (*DSP*). (b) Desmosome mutation frequency for each of 32 tumor types, with mutations aggregated across the 15 desmosome genes. Inset: Mutation frequency of each desmosome gene in skin cutaneous melanoma (SKCM). Other cancer type abbreviations: ACC - Adrenocortical carcinoma (N = 92), BLCA - Bladder urothelial carcinoma (N = 413), BRCA - Breast invasive carcinoma (N = 1108), CESC - Cervical squamous cell carcinoma and endocervical adenocarcinoma (N = 310), CHOL - Cholangiocarcinoma (N = 51), COADREAD - Colorectal adenocarcinoma (N = 640), DLBC - Lymphoid neoplasm diffuse large B cell lymphoma (N = 48), ESCA - Esophageal carcinoma (N = 186), GBM - Glioblastoma multiforme (N = 619), HNSC - Head and neck squamous cell carcinoma (N = 530), KICH - Kidney chromophobe (N = 113), KIRC - Kidney renal clear cell carcinoma (N = 538), KIRP - Kidney renal papillary cell carcinoma (N = 293), LAML - Acute myeloid leukemia (N = 200), LGG - Brain lower grade glioma (N = 530), LIHC - Liver hepatocellular carcinoma (N = 442), LUAD – Lung adenocarcinoma (N = 586), LUSC - Lung squamous cell carcinoma (N = 511), MESO - Mesothelioma (N = 87), OV - Ovarian serous cystadenocarcinoma (N = 617), PAAD - Pancreatic adenocarcinoma (N = 186), PCPG - Pheochromocytoma and paraganglioma (N = 184), PRAD - Prostate adenocarcinoma (N = 501), SARC - Sarcoma (N = 265), STAD - Stomach adenocarcinoma (N = 478), TGCT - Testicular germ cell tumors (N = 156), THCA - Thyroid carcinoma (N = 516), THYM - Thymoma (N = 124), UCEC - Uterine corpus endometrial carcinoma (N = 549), UCS - Uterine carcinosarcoma (N = 57), UVM - Uveal melanoma (N = 80). (c) Excess number of tumors with a mutated desmosome gene (colored points) above the number expected by chance (Online Methods). Horizontal black line indicates median value across desmosomal genes. Top ten most frequently mutated tumor types from (b) are shown. Significance determined using bootstrap analysis (*, q-value<10^−10^, Benjamani-Hochberg correction for multiple testing). (d) Ranked SKCM mutation frequencies of each human gene. Desmosome genes, red; known SKCM driver genes^22^, blue. PERP gene not shown in figure and is located at 63.5 percentile.

While these studies have focused on desmosome expression, a separate question is whether the desmosome is impacted by genetic mutations. Recently, we performed an integrative analysis of protein biophysical interactions and tumor mutations^17^, resulting in a map of 395 protein complexes under mutational selection in one or more cancer types. One of these complexes pointed to the desmosome as a potential focal point for accumulation of mutations in melanoma; however, at the time we did not validate or further explore this observation (Supplementary Fig. 1, Supplementary Table 1).

## Results

Here, to investigate the importance of mutations to the desmosome complex, we began with a comprehensive survey of somatic non-synonymous mutations impacting any of the 15 desmosome genes in aggregate, across 32 tumor tissue types profiled by The Cancer Genome Atlas (TCGA, Supplementary Table 2). By far the highest mutation recurrence was observed for cutaneous melanoma (SKCM), in which 71% of tumors harbored a desmosome coding mutation. In contrast, desmosome mutation rates were substantially lower in all other tissue types (<40%, Fig. 1b).

Since melanomas are one of the tumor types with generally high genome-wide burden of mutations^18^, we sought to assess the significance of desmosome mutation rates in several ways. First, we computed the excess mutational load on the 15 desmosome genes using MutSigCV^18^, which takes into account the genome-wide mutation burden, the type of mutational signature(s)^19^, and gene-specific factors such as mRNA expression level and time of replication during cell cycle. We found that as a group, desmosome genes are mutated significantly more often than expected, again with the strongest signal in melanoma (Fig. 1c). Second, we repeated this analysis in an independent melanoma cohort^20^, again finding that desmosome genes are highly and significantly mutated (Supplementary Fig. 2). These patterns did not depend on the particular mutational model for expected mutational frequency (P < 10^−20^ using MutSigCV^18^ and P < 10^−9^ using oncodriveFML^21^). Third, ranking human genes by decreasing mutation frequency in melanoma, we found that nearly all desmosome genes were among the top quartile, on par with the distribution of well-established melanoma cancer genes (Fig. 1d; with single exception of PERP). Notably, none of the desmosome genes had been previously reported to be significantly mutated in melanoma when analyzed individually^18,20,22^, perhaps since the mutational signal is spread across the desmosome complex.

The finding that the desmosome is frequently mutated in melanoma was puzzling, since in normal epidermis, desmosome function has been typically associated with keratinocytes rather than melanocytes^23^. However, this is a point of some confusion, since melanocytes cultured in the absence of other cell types have been shown to express the desmosomal cadherin Dsg2^24,25^ leading to increased migration^26^. To more fully understand the patterns and cell types associated with desmosome expression in melanoma tumors, we performed single-cell RNA sequencing (scRNA-seq via the inDrop system) on two human primary melanoma tumor biopsies encapsulating approximately 1,000 cells from each tumor^27,28^ (Fig. 2a). Single-cell transcriptomes were displayed using UMAP^29^ (Uniform Manifold Approximation and Projection) resulting in clusters that matched pre-identified distinct cell types (Online Methods). Analysis against a panel of established RNA markers for different human cell types distinguished clear populations of immune, epithelial, muscle, keratinocyte and melanocyte tumor types (Fig. 2b). Examining the expression profile of desmosome genes across these different types, we found desmosomes to be predominantly expressed in keratinocytes (Fig. 2c-d). To extend this analysis to animal models, we analyzed primary transgenic zebrafish tumors^30^, where we also found that desmosome expression is limited to keratinocytes (Supplementary Fig. 3a,b) as we had seen in the human melanoma samples. These results were also seen in normal (non-melanoma) skin cells, as indicated by analysis of a large existing single-cell RNA-Seq dataset from normal skin gathered by the Human Cell Atlas^31^, again showing that desmosomal gene expression is overwhelmingly restricted to keratinocyte cell populations (Supplementary Fig. 3c-e).

**Fig. 2:**
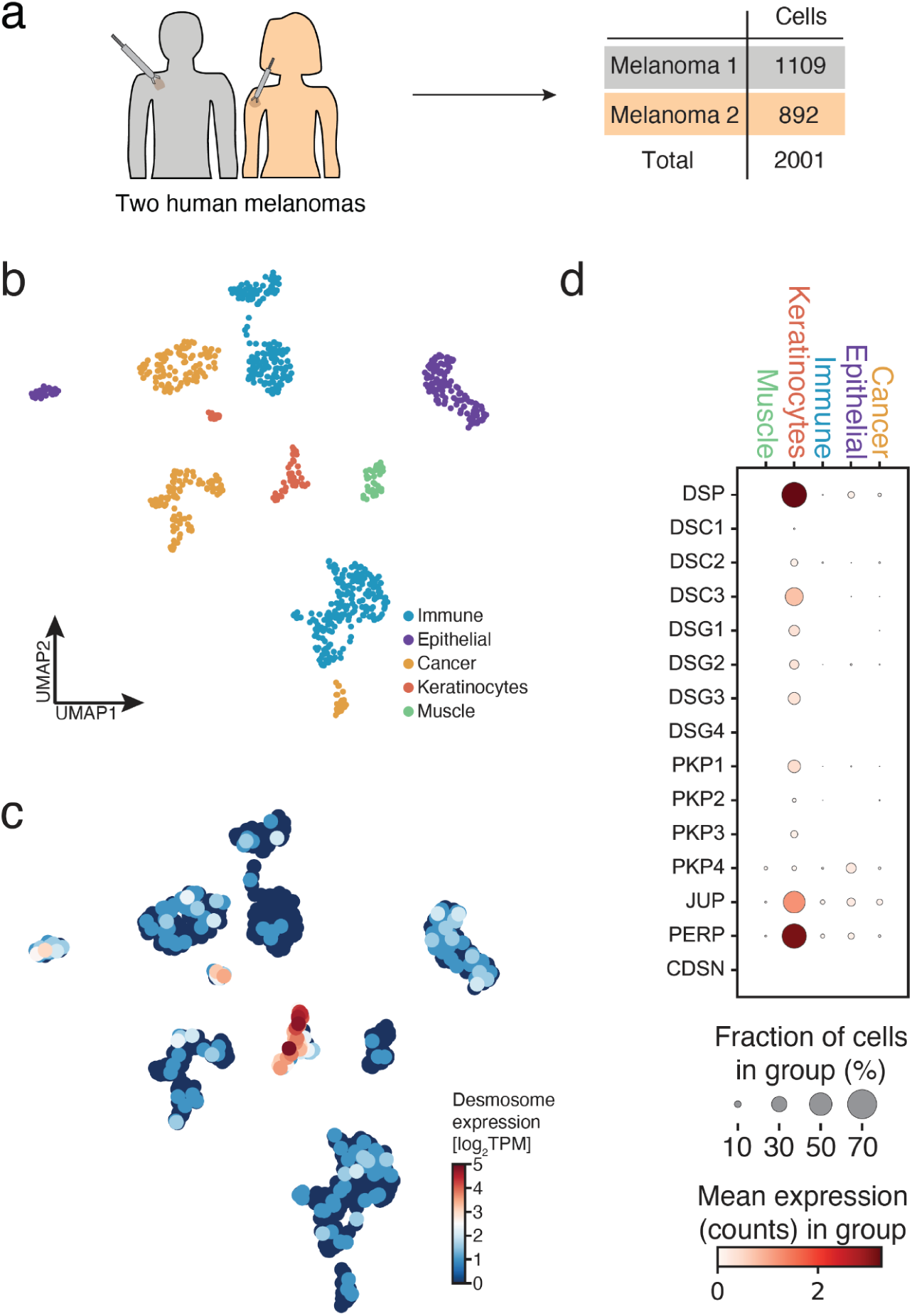
Single-cell analysis of desmosome gene expression in human melanomas. (a) Tumor biopsies from two patients, one male and one female, processed using scRNA-Seq. (b) Uniform Manifold Approximation and Projection (UMAP) analysis of 1109 individual cells from Melanoma 1, with color indicating inferred cell type. “Cancer” denotes melanoma cells. The two clusters of keratinocyte cells (red) reflect basal versus non-basal types. (c) Same UMAP as (b), with color indicating the expression of desmosome genes in transcripts per median (TPM, Online Methods). (d) Correspondence of desmosome gene expression to cell types. Color indicates mean expression per cell (log_2_). Point size indicates fraction of cells in cell type expressing each desmosome gene.

To investigate the relationship between desmosome expression and mutation, we next examined multi-omics data collected from 472 melanoma tumors by TCGA^32^ (104 primary and 368 metastatic, see Supplementary Fig. 4 for details about the SKCM cohort analyzed). Analysis of these data indicated that the expression of the desmosome system was significantly higher in primary than metastatic tumors, consistent with an elevated keratinocyte presence in primary versus metastatic melanoma (Fig. 3a). Examining the matched somatic mutation data, we found that primary tumors with desmosome mutations had significantly reduced expression of the desmosome complex in comparison to tumors without such mutations (Fig. 3b). Moreover, in primary melanoma only, there was a strong inverse relationship between the number of desmosome genes that are mutated in a tumor and overall desmosome expression (Fig. 3c). The most pronounced negative correlation between desmosome expression and desmosome mutations occurred in the early stages of melanoma, specifically in stage I and II tumors (Supplementary Fig. 5). Further analysis of this TCGA cohort found that desmosome mutation in primary melanoma associates with significantly higher expression of genes related to cell proliferation (Fig. 3d). To determine if these particular mutations are under evolutionary selection pressure, we calculated the ratio of nonsynonymous to synonymous mutations (dN/dS)^33^ in the desmosome genes. We found that dN/dS was approximately 2.5 in primary melanoma tumors, indicating strong positive selection, and that this value was significantly larger than for metastatic melanoma tumors (Fig. 3e). Finally, desmosome mutation was associated with a marginal increase in patient overall survival in this TCGA cohort (P = 0.03, FDR = 0.29; Supplementary Table 2). Collectively, these results suggested that accumulation of mutations in the desmosome disrupt its expression in a graded manner, and that these events are selected in primary melanoma.

**Fig. 3:**
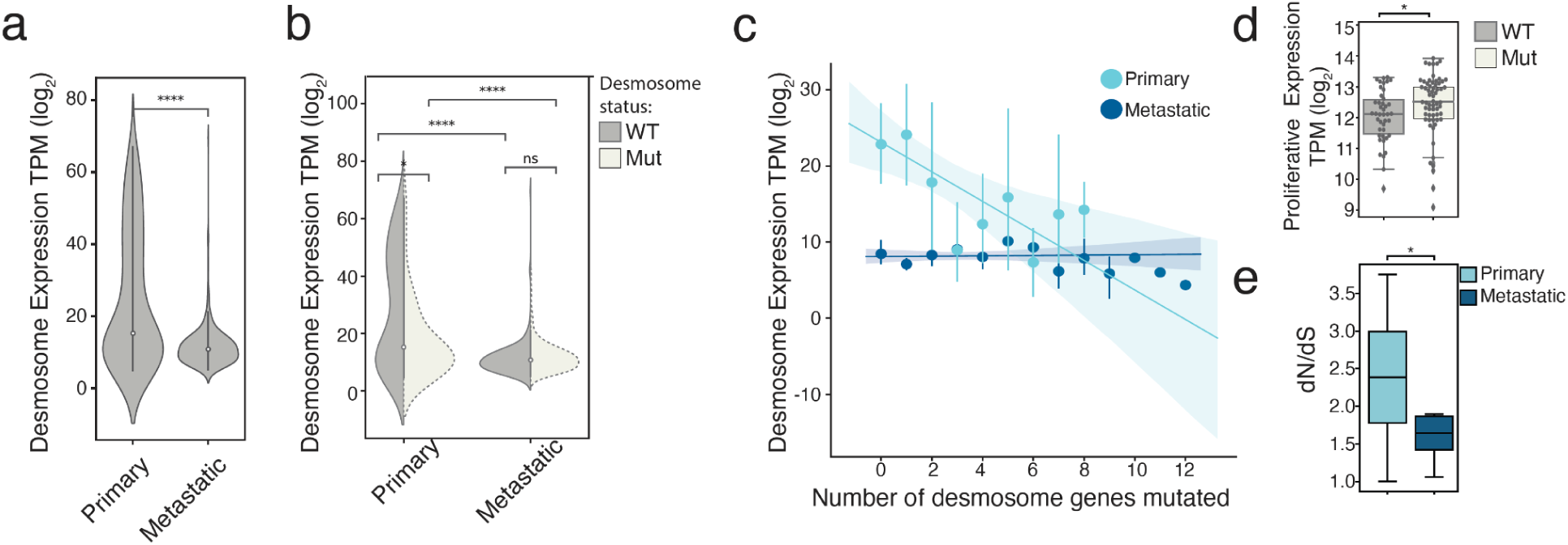
Relationship of desmosome expression to tumor grade and mutational status. (a) Violin plot of average desmosome gene expression in primary (left, N = 104) or metastatic (right, N = 362) melanoma tumors in TCGA. Significance determined by two sample t-test (****, P<10^−4^). TPM, transcripts per median. (b) Same as (a) distinguishing wild-type (WT, gray) and desmosome-mutated (white, N = 62 for primary tumors, N = 277 for metastatic tumors) tumors in the TCGA cohort. Significance determined by two sample t-test (****, P<10^−4^; *, P<0.05; ns, not significant). (c) Average desmosome gene expression (TPM) as a function of the number of desmosome genes with coding mutations. Shown separately for primary (light blue) versus metastatic (dark blue) melanoma tumor populations. Regression lines for each of these populations are shown: Primary melanoma, slope = –0.66; metastatic melanoma, slope = 0.03; significant difference with P<0.01. Error bars represent standard deviation and the shaded areas capture 95% confidence intervals. (d) Degree of proliferative gene expression (defined previously^70^) in desmosome WT (gray, N = 42) or desmosome mutant (white, N = 62) primary melanoma tumors (points) in the TCGA cohort. Significance determined by two sample t-test (*, P<0.05). The boxes in the plot contain the 25th to 75th percentile, the middle line denotes the 50th percentile, and the whiskers mark the 5th and 95th percentiles. (e) Ratio of fixed nonsynonymous-to-synonymous mutations (dN/dS) in primary (light blue, N = 12 genes across 104 patients) or metastatic (dark blue, N = 12 genes across 362 patients) melanoma tumors in the TCGA cohort. The distributions of dN/dS across the population of tumors are shown by box plots, using similar display convention to panel d. Significance determined by two sample t-test (*, P<0.05).

We reasoned that the distinction between primary and metastatic melanoma could be due to at least two explanations. First was the very different microenvironments of these two tumor locales. Primary melanoma tumor biopsies tend to contain large numbers of surrounding keratinocytes due to their growth within and below the epidermis, in contrast to metastatic tumors which, apart from cutaneous metastases, lack the epidermal component and are typically purer^34^. By this reasoning, the decrease in desmosome expression could be due to loss of desmosomal expression in keratinocytes. A second explanation was related to cancer-specific transcriptional states of melanoma cells. Since melanoma tumors can assume distinct transcriptional programs^35–37^, the effect on desmosome expression could be due to a change in transcriptional state of melanoma cells between primary and metastatic cancer.

To distinguish between these two possibilities, we performed spatial transcriptomics and protein immunofluorescence analysis of primary tumor biopsies drawn from a cohort of 18 melanoma patients (Fig. 4a). All samples were characterized by bulk whole-exome sequencing, identifying 9 tumors with desmosome mutations and 9 without (Supplementary Table 3). The first 8 samples were analyzed by NanoString GeoMx spatial transcriptomics profiling^38^, resulting in a deep mRNA expression profile for 4-8 regions of interest (ROI) per tumor. In each case, ROIs were selected to represent melanoma cells, keratinocytes or a combination of both cell types (Fig. 4b). Analysis of these data showed that ROI expression profiles were well-clustered according to cell type and desmosome expression levels (Supplementary Fig. 6). Desmosome expression was very high in keratinocyte-enriched ROIs but barely detectable in melanocyte-enriched ROIs (Fig. 4c), corroborating our earlier findings with scRNA-seq (Fig. 2b-c, Supplementary Fig. 3). Moreover, desmosome expression in keratinocyte ROIs was significantly lower in tumors with desmosome mutations than in tumors lacking such mutations (Fig. 4d, P<0.05). In contrast, we did not see substantial change in desmosome expression in melanocyte ROIs when comparing mutated versus unmutated samples (Fig. 4d). Finally, we found that tumors with desmosome mutations had significantly increased expression of numerous genes used as standard markers of cell proliferation, but only in ROIs enriched for melanocytes (Supplementary Fig. 7, Supplementary Table 4).

**Fig. 4:**
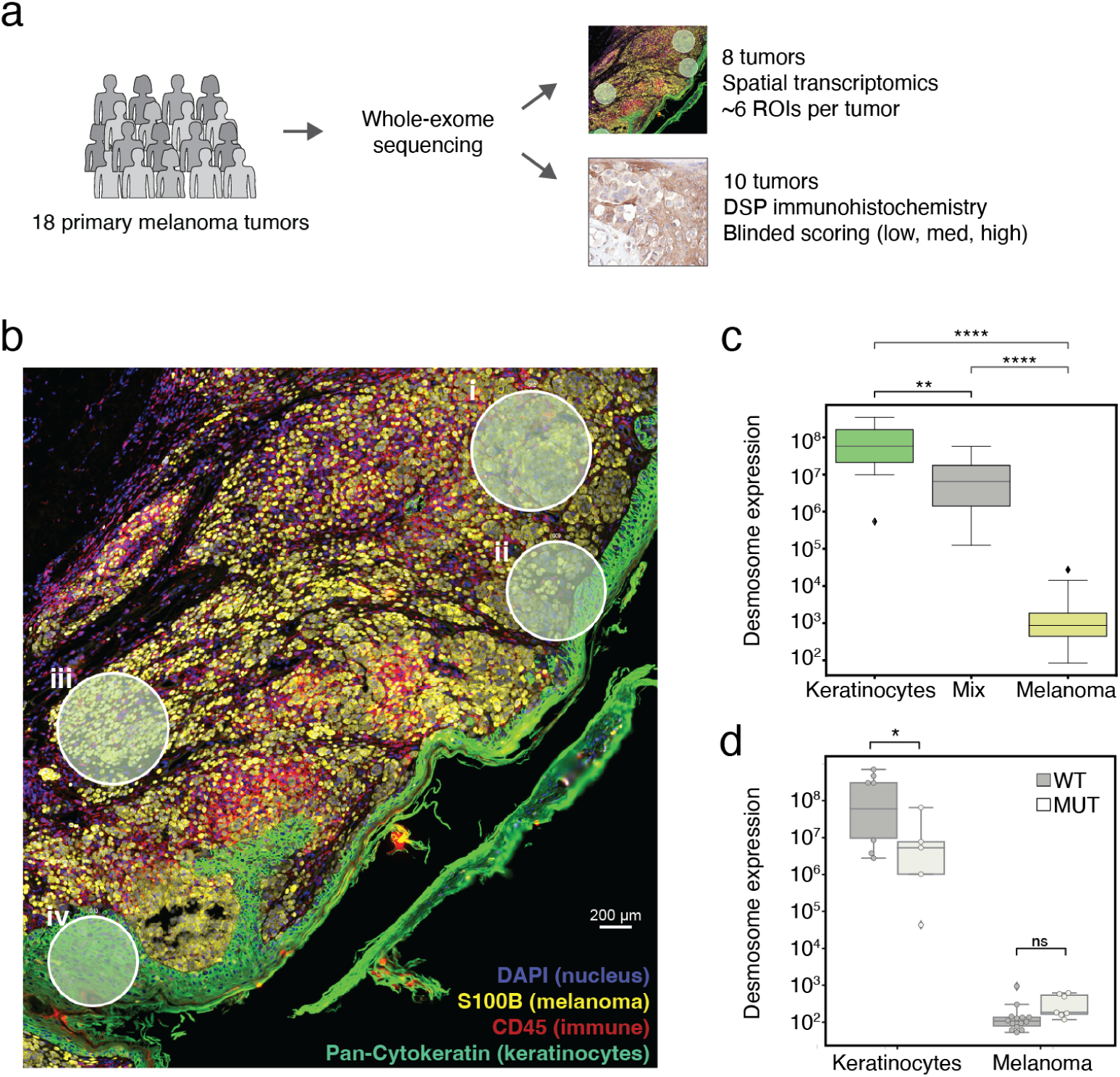
Spatial transcriptomics of tumors with mutated or wild-type desmosomes. (a) A total of 18 primary melanoma tumors underwent whole exome sequencing. Eight tumors were subjected to spatial transcriptomic analysis by NanoString GeoMx (top), covering a combined total of 48 regions of interest (ROIs). The remaining 10 tumors were subjected to immunohistochemistry (IHC) analysis in which expression levels of the DSP protein were scored by a dermatopathologist. The dermatopathology evaluation was performed in a blinded manner, without prior knowledge of the genomic status of desmosome mutations. (b) Tissue cross-section with example ROIs shown. Each ROI is indicated by a circle boundary marked by lowercase Roman numeral. ROIs are enriched for melanoma cells (i,iii), mixed melanoma cells and keratinocytes (ii), or keratinocytes (iv). Fluorescent markers label nuclei (blue), cancer cells (yellow), immune cells (red), or keratinocytes (green). (c) Average mRNA expression levels of desmosome genes in each ROI, stratified by keratinocyte versus melanoma ROIs. In the boxplots, lower and upper box boundaries delineate the 25th and 75th percentiles, the middle line denotes the 50th percentile, and whiskers mark the 5th and 95th percentiles. Significance determined by two sample t-test (**** P<10^−4^, ** P<10^−2^). (d) Average mRNA expression levels of desmosome genes in each ROI, stratified by keratinocyte versus melanoma and further subdivided based on whether ROIs are from tumors with wild-type (WT, dark gray) or mutant desmosomes (light gray). Within each category the set of ROI values is summarized by box plot, following the convention of panel c. Significance determined by two sample t-test (*, P<0.05, ns, not significant).

The remaining ten samples were used to perform immunofluorescence (IF) staining with an antibody against desmoplakin (DSP), the central desmosomal protein which provides links to keratin intermediate filaments in the cytosol (Fig. 5a). DSP had been shown previously to be essential to desmosome structural integrity^39^, and we had also found it had the single largest mutation burden in melanoma patients (Fig. 1c-d). In addition to DSP, the samples were co-stained with antibodies against SOX10 (melanocyte nucleocytoplasmic marker) and JUP (keratinocyte membrane marker). The analysis revealed that samples harboring desmosome mutations exhibited DSP expression levels that were significantly lower than DSP levels in samples without such mutations (Fig. 5b). This difference was significant not only for total DSP intensity but also when quantifying DSP intensity at the cell membrane (Fig. 5b), as the majority of DSP was specific to that location (Fig. 5a). Thus, both the earlier spatial profiling (Fig. 4) and the subsequent immunofluorescence data (Fig. 5) supported a model in which desmosome mutation leads to a reduction in desmosome expression specifically in keratinocytes.

**Fig. 5.**
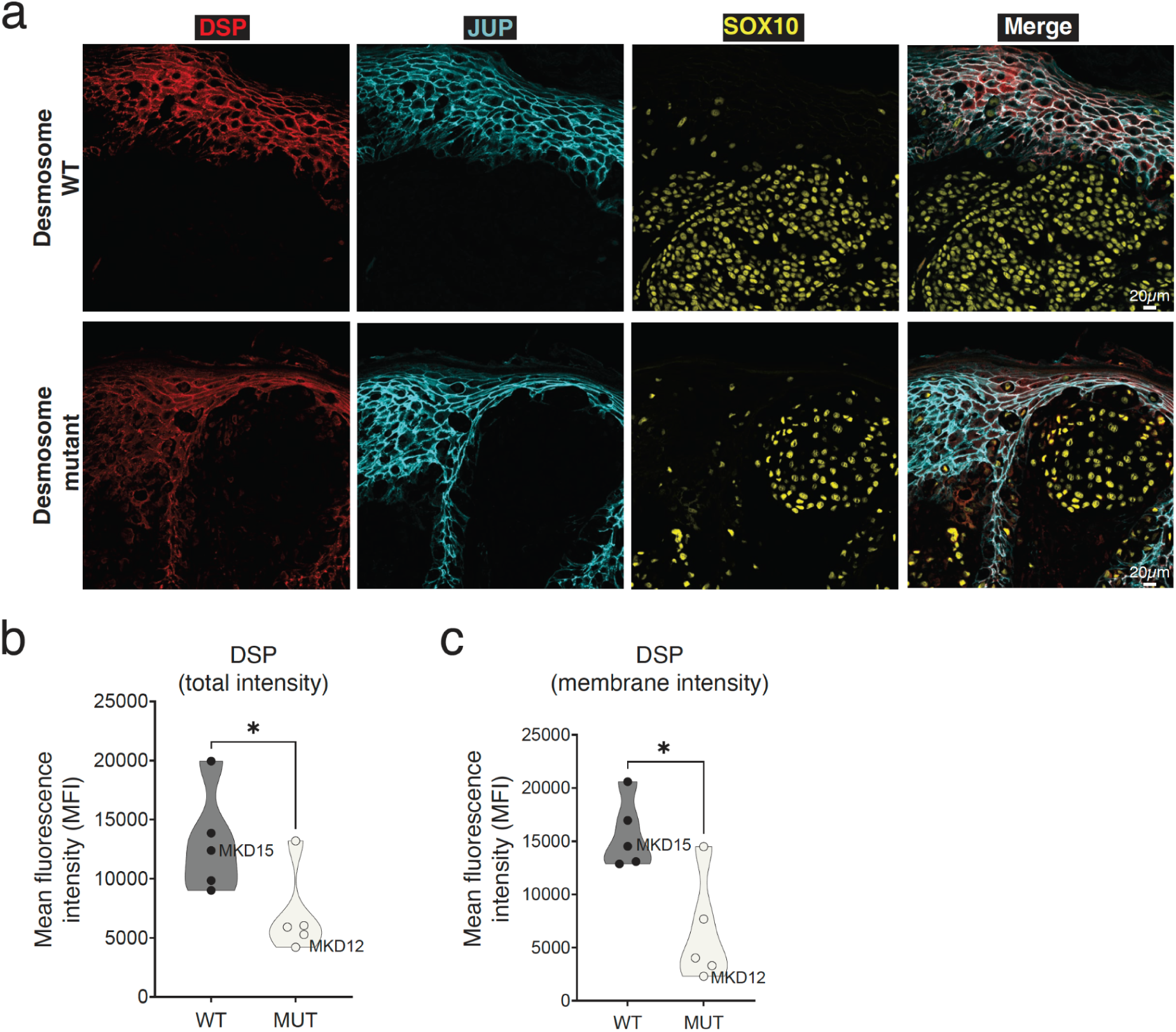
DSP Immunofluorescence of tumors with mutated or wild-type desmosomes. (a) DSP Immunofluorescence of tumor MKD15 (top, desmosome wild-type) and MKD12 (bottom, desmosome mutant). Red color indicates DSP expression, cyan color represents JUP expression serving as a membrane mask, yellow color represents SOX10 expression marking melanocytes. (b) Total DSP protein expression levels across 10 human melanoma tumor adjacent epidermis primary tumors. Samples are split by desmosome mutational status (WT, MUT). Significance by Wilcoxon rank-sum test (*, P<0.05). (c) Same as (b) but for membrane-only DSP protein expression.

We next wanted to functionally test whether loss of desmosome expression in keratinocytes can influence the behavior of neighboring melanoma cells. For this purpose, we used a keratinocyte cell line (HaCaT human immortalized keratinocytes, Online Methods), which we demonstrated reliably expresses and localizes the desmosome at cell-cell junctions, with the expected connections to cytokeratins (Fig. 6a). We then subjected this cell line to stable knockdown of *DSP* gene expression using shRNA (Supplementary Table 5), which we confirmed causes marked loss of DSP protein along with the loss of other desmosome components such as DSG2 as reported earlier^40^ (Fig. 6b, Supplementary Fig. 8). Next we examined effects of DSP knockdown on cell proliferation in a series of complementary cell culture experiments, including keratinocyte and melanoma monocultures, keratinocyte/melanoma co-cultures, and keratinocyte-to-melanoma conditioned media transfer assays (Fig. 7a). In monocultures, *DSP* knockdown led to no change in proliferation in the HaCaT keratinocytes or melanoma cells over several days in culture (Supplementary Fig. 9a). In addition, *DSP* knockdown did not result in any change in apoptosis in melanoma cells and keratinocytes (Supplementary Fig. 9b-c). In contrast, when keratinocytes and melanoma cells were grown together in co-culture, we found that proliferation of melanoma cells increased by more than two-fold upon *DSP* knockdown in keratinocytes (Fig. 7b, Supplementary Fig. 9a). Notably, significant increases in melanoma proliferation could also be obtained by transfer of conditioned media from *DSP*-knockdown keratinocytes to melanoma monocultures (Fig. 7c); this effect was seen in three independent melanoma cell lines, two of which are derived from primary melanoma tumors in the radial growth phase (RGP)^41,42^. These results suggested that loss of keratinocyte desmosomes promotes neighboring melanoma growth through secreted factors (Fig. 7d).

**Fig. 6:**
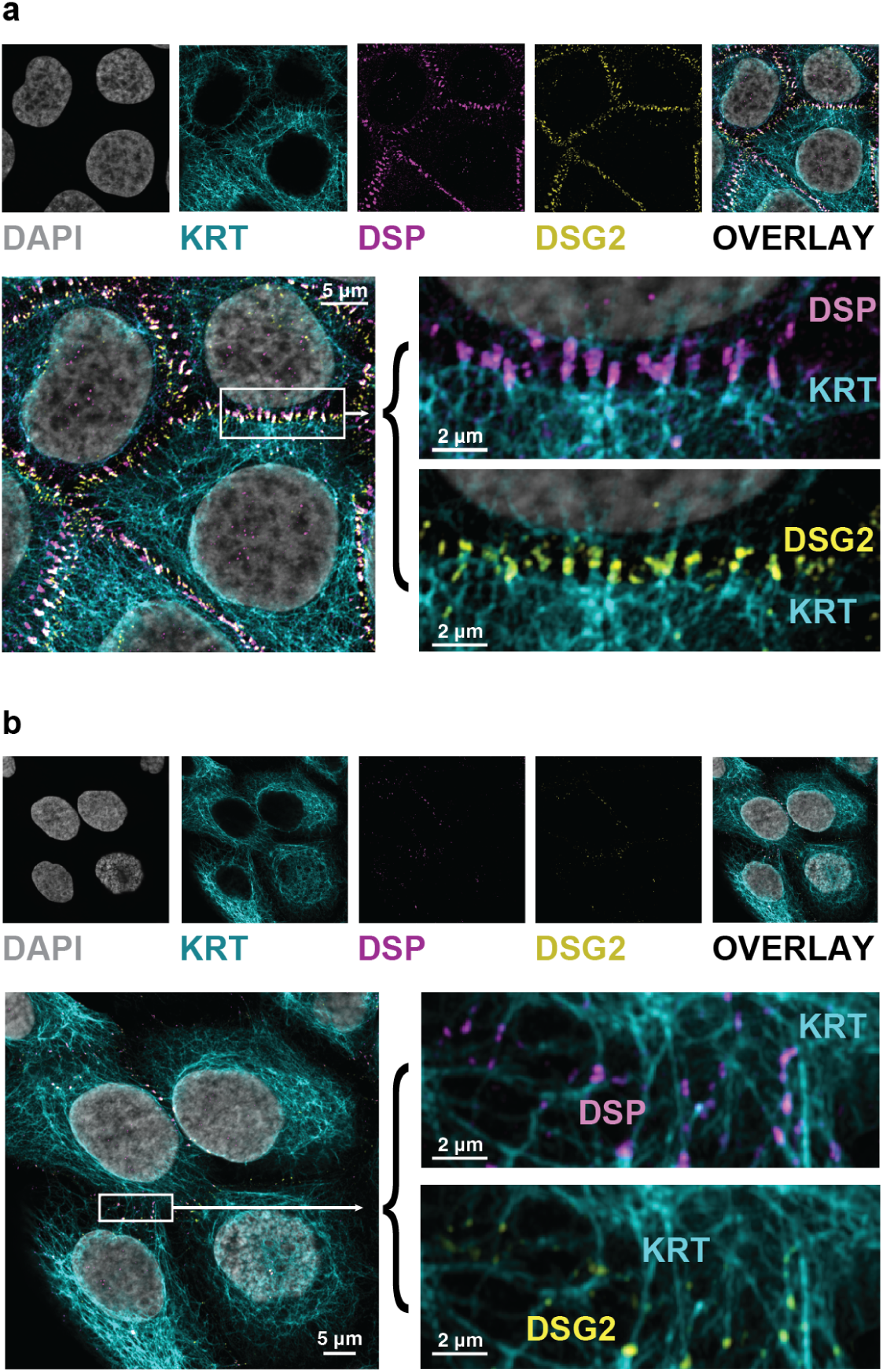
Visualization of desmosomes at keratinocyte cell junctions –/+ DSP knockdown. (a) Representative image from keratinocyte monocultures expressing stable non-targeting control (NTC) shRNA, depicting staining for keratin (KRT14, cyan), desmoplakin (DSP, purple), and desmoglein (DSG2, yellow). Cell nuclei stained with DAPI (gray). Each stain is shown separately (upper small images) as well as in a single image overlay (right-most upper image and lower enlargements). The images show robust expression and expected location of desmosome components (DSP, DSG2). (b) Same as panel (a) but for keratinocyte monocultures expressing stable DSP shRNA, illustrating effective knockdown of desmosome proteins DSP and DSG2.

**Fig. 7:**
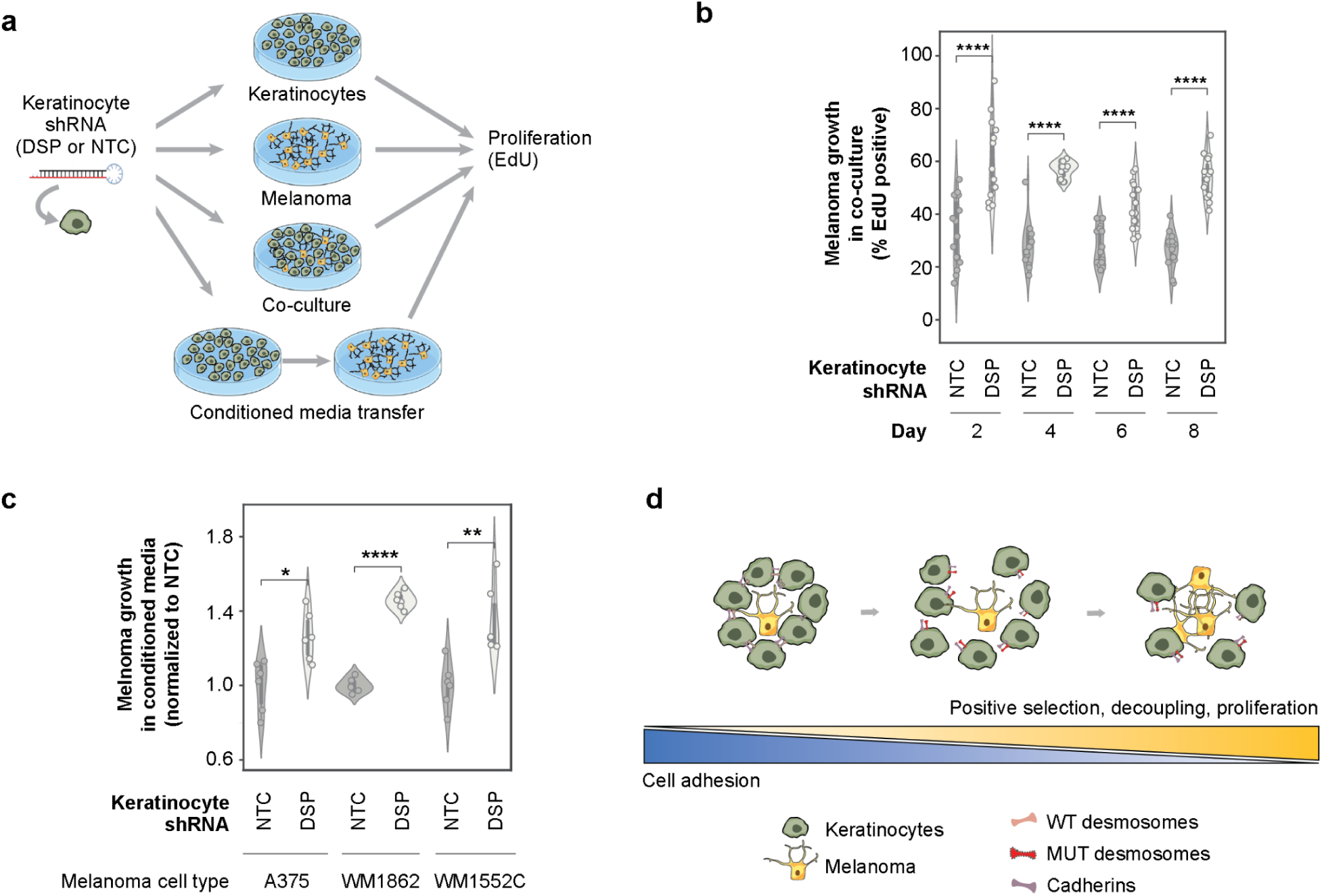
Effects of *DSP* gene disruption in melanoma/keratinocyte monocultures and co-cultures. (a) Keratinocytes were engineered to stably express either non-targeting control (NTC) shRNA or *DSP* shRNA constructs; melanoma cells were not genetically engineered. These cell types were grown and characterized independently or as co-cultures (first three rows). Alternatively, melanoma cells were characterized in keratinocyte-conditioned media (final row). Cells were stained with specific antibodies and characterized by proliferation assays. (b) EdU assay of melanoma cell proliferation (A375), shown for keratinocyte/melanocyte co-cultures in which keratinocytes stably express NTC shRNA (dark gray) versus *DSP* shRNA (light gray) for 2, 4, 6 or 8 days. Each violin summarizes 5 distinct fields per well ⨉ 3 biological replicates for N = 15 replicate measurements. (****, P<0.0001) (c) Cyquant assay of melanoma cell proliferation when grown in conditioned media from keratinocytes stably expressing either NTC shRNA (dark gray) or *DSP* shRNA (light gray). Results for three different melanoma cell lines are shown (A375, metastatic; WM1862 and WM1552C, non-metastatic). Each violin summarizes six conditioned-media measurements. (* P<0.05, ** P<0.01, **** P<0.0001) (d) Suggested model in which desmosome mutations in keratinocytes lead to decoupling of keratinocyte-melanoma cells and decreased keratinocyte adhesion which, in turn, promotes melanocyte proliferation.

## Discussion

Typically, studies of the cancer genome have focused on identifying frequent genetic alterations in single genes, or associations between individual gene alterations and tumor functional outcomes^43–46^. The main challenge of these gene-by-gene approaches is that most cancer mutations are rare. However, consideration of groups of genes, corresponding to discrete cellular components such as protein complexes and signaling pathways, provides an expanded means of understanding mutational effects^17,47–49^. Using such a strategy we have shown that the desmosome complex is frequently impacted by coding mutations in primary melanoma samples, associated with a substantial decrease in expression of desmosome components. Spatial transcriptomic profiling and protein immunofluorescence indicate these expression decreases occur predominantly in keratinocytes in the tumor local microenvironment rather than in melanoma tumor cells. These observations are corroborated by genetic disruptions showing that loss of the DSP desmosome subunit in keratinocytes has the effect of amplifying the proliferation of adjacent melanoma cells in co-culture.

Our finding that some tumors harbor coding mutations in several desmosome genes, not just one (Fig. 3c, Supplementary Fig. 4), may relate to several underlying reasons. Some genes in the desmosome system are paralogs, for example the cadherins DSG1, DSG2 and DSG3, and studies have shown that loss of DSG3 function can be compensated by DSG2^50,51^. Similarly, desmocollins can undergo expression pattern switching, in which loss of expression of one isoform can cause upregulation of alternative isoforms thus retaining desmosome activity^52^. It is thus possible that loss of certain desmosome genes does not provide a selective advantage in melanoma due to functional redundancy with alternate gene family members.

We also observed differences in desmosome mutation patterns across melanoma cohorts (Fig. 1c, Supplementary Fig. 2). While these may relate to underlying differences in the biology of the tumor populations analyzed (TCGA-SKCM versus Conway et al.), it is clear that at least part of the difference can be attributed to variations in the two data processing pipelines. In particular, Conway et al. describe a more stringent process than that of the TCGA-SKCM study, for instance using an “expression filter” based on a human single-cell RNA-Seq dataset among other criteria. Despite these differences, our system-level approach was able to identify the desmosome system as significantly mutated in both cohorts.

The observed interplay between keratinocytes and melanoma cells stands in contrast to the usual mode by which somatic mutations are thought to promote cancer, by direct selection in a clonal population of tumor cells. It is unclear whether desmosome-mutant keratinocytes are actually selected by the melanoma tumor, or merely serve as a pre-existing condition that increases risk of tumor formation. A recent study using organotypic skin cultures highlighted that melanoma cells can transcriptionally repress the expression of a desmosomal cadherin subunit, DSG1, in keratinocytes, via paracrine signaling^53^. Further, loss of DSG1 expression correlates with increased migration of melanoma tumor cells, suggesting that downregulation of desmosome gene expression is pro-tumorigenic in melanoma, independent of mutations^53^. If selection does occur, one mechanism could be that desmosome-mutant keratinocytes and proliferating melanocytes synergistically amplify one another’s growth. We do not yet know how “large” a patch of mutant keratinocytes would need to be to affect a nearby melanoma. Regardless of clone size, our data suggest that the keratinocyte-melanoma interaction may occur through secreted factors, the nature of which will need to be identified in future studies.

While the mRNA expression, protein immunofluorescence, and functional cell culture experiments suggest that desmosome mutations are primarily important in keratinocytes (Fig. 4d, Fig. 7b-c), a limitation of our study is that we have not yet conclusively shown that such mutations are specifically enriched in keratinocytes compared to melanoma cells. The reasons for this limitation are largely technical, as single-cell and spatial DNA sequencing are far less developed than comparable technologies for RNA. In addition, primary melanomas (most relevant for our study) are typically very small with little tissue left over for experimental work. As single-cell DNA technologies improve, they will undoubtedly shed important light on our findings. Very relevant to this question is a recent preprint describing a process for whole-exome sequencing of single cell clonally-expanded normal skin cells^54^. A cursory analysis of these clonal DNA sequencing data shows accumulation of desmosome mutations in both keratinocytes and melanocytes (Supplementary Fig. 11a,b), with the highest number of unexpected mutations falling within the DSP gene in keratinocytes (Supplementary Fig. 11c). These single-cell mutation data are not from melanoma tumors, however, and are still of small sample size.

More generally, our results are consistent with earlier findings that non-tumor cells can indeed carry cancer-causing mutations, predominantly in genes associated with tumor initiation rather than clonal expansion^55–57^. For example, a study of melanocyte transformation identified that non-tumor cells close to a melanoma tumor have significantly greater numbers of mutations than those farther away, including mutations in oncogenes such as *BRAF*^58^. Our findings not only support the notion that tumors arise from a collection of cells rather than a clonal population^59–62^, but they extend that notion to other cell types which are not necessarily the cell type of origin. By such a model, mutations in complementary cellular communities may alter expression programs and growth profiles which, in turn, help determine the order in which specific mutations arise within the complex cancer tissue.

## Online Methods

### Statistical framework for assessing desmosome mutation

The enrichment analysis of coding mutations within the desmosome system was conducted for each TCGA cancer cohort following the method of Zheng et al.^17^. First, we used the MutSigCV^18^ tool to estimate the expected number of tumor samples with somatic coding mutations in each human gene, assuming no positive selection pressure. In estimating this expected number for each gene, MutSigCV accounts for an array of features including gene length, local mutational signatures, DNA replication timing, and mRNA expression level. Second, for each gene we computed the difference between the actual and expected counts of mutated tumor samples. Genes were ranked by decreasing value of this difference in an ordered gene list. Third, this list was examined for the significance of the rankings of desmosome genes using permutation testing (similar to Mann Whitney U Test). For the permutation test, the average rank of the desmosome genes in the ordered list was compared to their average rank in each of 100,000 permuted lists of all human genes. This test generated a p-value for each TCGA cancer cohort. To account for multiple hypothesis testing (i.e. due to multiple cohorts), p-values were adjusted using the Benjamini-Hochberg procedure, resulting in q-values.

### Single-cell RNA sequencing of primary melanoma tumors

Melanoma tumors were collected post-operatively from two patients who consented and signed 604 IRB. To obtain single-cell suspensions, samples were washed in PBS and cut into small pieces (4-5 mm^3^) followed by dissociation using the Miltenyi human tumor dissociation kit according to manufacturer instructions. Red blood cells were depleted using ACK lysis buffer for 3 minutes. Dead cells were removed, if needed, using the Milyenty dead cell separator. Single cell encapsulation and library preparation were performed using the inDrop platform^27,28^ followed by sequencing on an Illumina NextSeq. Sequencing reads were de-multiplexed, aligned and counted using a custom pipeline as described previously^36^ to produce a raw count matrix for each tumor. Quality control was also performed as described previously^63^. The raw matrix count was first transformed to UMIs (Unique Molecular Identifiers) to correct for library preparation duplications^64^. Cells with >750 distinct UMIs, mitochondrial transcripts <20% and ribosomal transcripts <30% were selected for analysis. UMI counts for the selected cells were normalized by the total number of transcripts per cell, and a scale factor equivalent to the median number of transcripts across all cells was applied (transcripts per median, TPM). Expression was transformed using the Freeman-Tukey transform as described previously^65^.

### UMAP 2D projection and annotation of single-cell mRNA-seq data

We selected genes with both above-mean mRNA expression level and above-mean Fano factor (a standardized measure of variance). These genes were then used as features to project cells onto two dimensions using the Uniform Manifold Approximation and Projection for Dimension Reduction (UMAP) technique with default parameters^29^. Ten communities of cells were clearly discernible in the resulting UMAP projection (Fig. 2b,c). To label the cell type represented by each community, we identified the top differentially expressed genes in each (p < 10^−6^, Kolmogorov Smirnov test; effect size > 0.2, Cohen’s d). These genes were cross-referenced against a collection of published gene expression markers of known cell types^30,35,37,66,67^, allowing unambiguous assignment of each community to one of five types: immune (markers: *CD45*, *CD4*, *CD8A*), epithelial (*EPCAM*), muscle (*MYH11*), melanocyte (*SOX10*, *SOX2*, *S100B*) or keratinocyte (*KRT4*, *KRT5*, *KRT17*, *KRT19*). To further annotate a subset of melanocytes as malignant, we inferred copy-number variations (CNV) from the RNA-seq profiles with inferCNV^68^, using all other cells from the same tumor sample as a reference. All three melanocyte communities were found to have a common pattern of aneuploidy in their CNV profiles, indicating cancer.

### Reanalysis of Single-Cell RNA Sequencing Data Single-cell RNA sequencing data from the Human Cell Atlas skin dataset

This dataset was reanalyzed using the CellAtlas.io application. The analysis focused on key relevant cell types, including melanocytes, undifferentiated keratinocytes, and differentiated keratinocytes. Out of the 15 desmosome genes evaluated, 11 were detected in the dataset (see Supplementary Fig. 3d-e).

### Spatial transcriptomics by GeoMX platform

Formalin-fixed paraffin-embedded (FFPE) samples of eight primary melanomas were chosen and sectioned (clinical information provided in Supplementary Table 2). Three sections were taken from each tumor: two were processed for whole-exome sequencing to determine desmosome mutation status, and one was retained for the digital spatial profiling workflow to determine desmosome expression. Digital spatial profiling was performed using the GeoMX system as previously described^69^. Immunofluorescent visualization markers for keratinocytes (*PanCK*), melanoma cells (*S100B*, *PMEL*) and immune cells (*CD45*) were used to guide the selection of regions of interest (ROIs) containing either pure or mixed areas of each cell type (keratinocyte, melanocyte) followed by RNA profiling using the GeoMx human whole assay.

### Cell lines (melanoma and keratinocytes)

Melanoma cell line A375 was obtained from ATCC (August 2013), authenticated by STR profiling and tested for *Mycoplasma* contamination using a MycoAlert *Mycoplasma* Detection kit (Lonza #: LT07-318). Authenticated and tested primary RGP melanoma cell lines WM1862 and WM1552C were obtained from Rockland Inc. (Dec 2023). A375 cells were maintained in DMEM (Gibco #11965) supplemented with 10% FBS (Gemini Bio) and 1X penicillin/streptomycin (Gibco #15140122). WM1862 and WM1552C were maintained in tumor-specialized medium with 2% FBS according to the manufacturer’s instructions. HaCaT (immortalized keratinocyte cell line) was obtained from Addexbio and authenticated by the MSKCC Molecular Cytogenetics Core. Keratinocyte lines were maintained in DMEM (Gibco #11965) supplemented with 10% FBS (Gemini Bio) and 1X penicillin/streptomycin.

### Lentiviral shRNA cell lines

For desmoplakin (DSP) knockdown studies we used Dharmacon SMARTVector lentiviral shRNA constructs (n = 3 distinct DSP-targeting shRNAs). Of these shRNAs, 2 showed >80% knockdown and were pooled for subsequent studies (Supplementary Table 5). Stable knockdown cell lines (HaCaT and A375) were created using puromycin selection for one week. HaCaT cells stably expressing NTC (Non-targeting control) shRNA or DSP targeting shRNA were then co-cultured with the indicated melanoma cell lines for the time points indicated. HaCaT and A375 cells in monoculture stably expressing NTC (Non-targeting control) shRNA or DSP targeting shRNA were subsequently plated in 96-well plates and used for plate-based proliferation assays.

### Immunostaining

For immunostaining, melanoma/keratinocyte monocultures or co-cultures were fixed in 4% PFA for 15 – 30 min @ RT followed by 3X PBS washes. Cells were permeabilized in 0.1% Triton-X for an additional 15 – 30 min, followed by 3X PBS washes. For the EdU stain, click-chemistry-based EdU detection was performed using an Abcam kit (#ab219801) according to manufacturer’s instructions. To stain for cell-specific markers, cells were treated overnight with primary antibodies at 4°C then subsequently washed with PBS and incubated with secondary antibodies and a nuclear stain (Invitrogen #H3570). Imaging was performed on an LSM880 high-resolution confocal microscope with a 63X objective using AiryScan imaging. The following primary antibodies were used: mouse anti-human KRT14 (Abcam #ab7800); rabbit anti-human SOX10 (ThermoFisher #PA5-84795); rabbit anti-human S100A6 conjugated to Alexa Fluor 647 (Abcam #ab204028); guinea pig anti-human desmoplakin (Progen #DP-1); mouse anti-human desmoglein 2 (ThermoFisher #32-6100). The percentage of EdU-positive cells was quantified by calculating double-positive cells (SOX10+/S100A6+ or KRT14+ and EdU+) as a fraction of the total number of SOX10/S100A6+ or KRT14+ cells in each field. Image analysis was performed using FIJI. For the DSP Immunofluorescence, tissue sections of 5-μm thickness were cut from formalin-fixed, paraffin-embedded tissue blocks. Immunostaining was performed using standard deparaffinization, heat induced epitope retrieval (HIER) procedures as described previously^3^. Immunostained slides were counterstained with Hoechst 33342 (Invitrogen #H3570) and mounted in ProLong Glass Antifade Mountant (Fisher Scientific #P36984). The following primary antibodies were used: anti-rabbit SOX10 (Cell Signaling E6B6I #69661), anti-mouse JUP (Santa Cruz, sc514115), anti-guinea pig DSP (Progen, DP-1). For DSP membrane signal intensity quantification, JUP staining was used as a membrane mask and quantifications were performed using ImageJ. All sections were imaged on an LSM880 high resolution confocal microscope with a 20X objective.

### Cyquant proliferation assay in monocultures

For plate-based measurement of proliferation, 2000 or 5000 cells/well (melanoma cells or keratinocytes) were plated in a 96-well plate followed by 24 – 96 hour incubation. Proliferation was measured using a Cyquant Cell Proliferation assay (Invitrogen #C7026) as per Manufacturer’s instructions. For conditioned media experiments, equal cell numbers of NTC or DSP shRNA-expressing keratinocytes were cultured in fresh medium; this medium was then collected after 48 hours and used to treat the melanoma cells. Fluorescence was ascertained using a BioTek Synergy 96-well plate reader, with all values normalized to the control conditions.

## Data Availability

All major ‘omics datasets relevant to this study will be deposited in relevant public databases upon acceptance of this article for publication. These datasets are:

| Dataset | Database | Accession Number |
| --- | --- | --- |
| Single-cell RNA sequencing of primary melanoma tumors | Gene Expression Omnibus | TBD |
| Spatial transcriptomics by GeoMX platform | Gene Expression Omnibus | TBD |

## Code Availability

All code used for this work, including custom algorithms and analysis scripts, has been distributed on GitHub at the URL: http://github.com/MaayanBaron/desmosome_paper.git.

## Supplementary and Extended Information

Supplementary Tables 1 - 5

Supplementary Figures 1 – 11

Extended Data Table 1 (Online)

## Author Contributions

M.B. and T.I. conceived and implemented the study, M.T., M.B. and R.W. designed and performed the functional work. D.B. M.B. and I.Y. performed and analyzed the single cell RNA-Seq work. M.B., T.I. and M.K. performed and analyzed the Spatial Transcriptomics work. F.Z., P.W. and J.Y. provided critical interpretation. The paper was written by M.B. and T.I. and was critically reviewed and approved by all of the coauthors.

## Declaration of Interests

T.I. is a co-founder, member of the advisory board, and has an equity interest in Data4Cure and Serinus Biosciences. T.I. is a consultant for and has an equity interest in Ideaya BioSciences. The terms of these arrangements have been reviewed and approved by UC San Diego in accordance with its conflict of interest policies. R.M.W. is a paid consultant to N-of-One, a subsidiary of Qiagen.

## Acknowledgements

We gratefully acknowledge funding support from the National Institutes of Health under grants NCI U54 CA274502 and NIGMS P41 GM103504 to TI and grant R03CA277645 to MK. We would also like to thank Lan Yu and Aubrey Gasper for technical assistance with histology and immunohistochemistry for this study and Stephenie Liu and Ryan Davis for assistance with whole exome sequencing, supported by the UC Davis Comprehensive Cancer Center Support Grant awarded by the National Cancer Institute (NCI P30CA093373).

## Supplementary Tables and Figures

**Supplementary Table 1:** Systems under mutational pressure^†^ in melanoma TCGA-SKCM.

| Rank | System name | System annotation | p-value | FDR |
| --- | --- | --- | --- | --- |
| 1 | Cluster4-315 | ECM_Collagen1 | 0.0001 | 0.0585 |
| 2 | Cluster7-18 | ECM_Collagen2 | 0.0001 | 0.0585 |
| 3 | Cluster9-1 | ECM_Collagen3 | 0.0001 | 0.0585 |
| 4 | Cluster3-231 | MAPK_cascade1 | 0.0001 | 0.0585 |
| 5 | Cluster4-90 | Cell_cycle1 | 0.0003 | 0.1403 |
| 6 | Cluster3-334 | Desmosome | 0.0011 | 0.2338 |
| 7 | Cluster2-374 | Transcription | 0.0011 | 0.2338 |
| 8 | Cluster7-155 | SWI/SNF | 0.0009 | 0.2338 |
| 9 | Cluster2-30 | MAPK_cascade2 | 0.0007 | 0.2338 |
| 10 | Cluster3-341 | E2F | 0.0006 | 0.2338 |
| 11 | Cluster6-233 | Cell_cycle2 | 0.0008 | 0.2338 |

<sup>†</sup> Mutated systems with FDR < 0.25 as determined in previous study, see Zheng et al.<sup>17</sup> for details.

**Supplementary Table 2:** TCGA cohorts and association of desmosome mutation with survival^†^.

| TCGA cohort | Overall Survival |  |  | Disease Free Interval |  |  | Total Patients | WT number of patients | MUT number of patients |
| --- | --- | --- | --- | --- | --- | --- | --- | --- | --- |
|  | p-value | FDR | Poor Prognosis direction | p-value | FDR | Poor Prognosis direction |  |  |  |
| ACC | 0.0019 | 0.0627 | MUT | 0.9451 | 1 | NA | 92 | 78 | 14 |
| THCA | 0.009 | 0.1478 | MUT | 0.3295 | 0.6739 | NA | 580 | 569 | 11 |
| SKCM | 0.0267 | 0.2937 | WT | NA | NA | NA | 472 | 137 | 335 |
| UCEC | 0.0641 | 0.4068 | NA | 0.0286 | 0.2359 | NA | 583 | 396 | 187 |
| KICH | 0.0643 | 0.4068 | NA | 0.6605 | 0.9 | NA | 91 | 86 | 5 |
| CHOL | 0.074 | 0.4068 | NA | 0.2693 | 0.6739 | NA | 45 | 42 | 3 |
| OV | 0.1485 | 0.6618 | NA | 0.4344 | 0.7168 | NA | 602 | 544 | 58 |
| SARC | 0.1774 | 0.6618 | NA | 0.8409 | 1 | NA | 271 | 248 | 23 |
| BLCA | 0.1805 | 0.6618 | NA | 0.3622 | 0.6739 | NA | 436 | 306 | 130 |
| PCPG | 0.2076 | 0.685 | NA | 0.8496 | 1 | NA | 187 | 184 | 3 |
| MESO | 0.2439 | 0.7316 | NA | 0.3429 | 0.6739 | NA | 87 | 81 | 6 |
| ESCA | 0.2666 | 0.7332 | NA | 0.6177 | 0.8862 | NA | 204 | 179 | 25 |
| UCS | 0.3248 | 0.8244 | NA | 0.3615 | 0.6739 | NA | 57 | 47 | 10 |
| PAAD | 0.4081 | 0.874 | NA | 0.0232 | 0.2359 | MUT | 196 | 189 | 7 |
| UVM | 0.4215 | 0.874 | NA | NA | NA | NA | 80 | 77 | 3 |
| LIHC | 0.4238 | 0.874 | NA | 0.1432 | 0.6739 | NA | 438 | 391 | 47 |
| KIRC | 0.5402 | 0.9426 | NA | 0.9394 | 1 | NA | 944 | 915 | 29 |
| PRAD | 0.5482 | 0.9426 | NA | 0.2857 | 0.6739 | NA | 566 | 546 | 20 |
| LGG | 0.5856 | 0.9426 | NA | 0.4275 | 0.7168 | NA | 529 | 477 | 52 |
| THYM | 0.5876 | 0.9426 | NA | NA | NA | NA | 126 | 120 | 6 |
| COAD | 0.6211 | 0.9426 | NA | 0.6818 | 0.9 | NA | 545 | 443 | 102 |
| DLBC | 0.6691 | 0.9426 | NA | 0.3412 | 0.6739 | NA | 48 | 37 | 11 |
| STAD | 0.6861 | 0.9426 | NA | 0.4824 | 0.7236 | NA | 511 | 394 | 117 |
| CESC | 0.6873 | 0.9426 | NA | 0.2479 | 0.6739 | NA | 312 | 250 | 62 |
| KIRP | 0.7603 | 0.9426 | NA | 0.9111 | 1 | NA | 352 | 316 | 36 |
| LUSC | 0.7866 | 0.9426 | NA | 0.3676 | 0.6739 | NA | 623 | 454 | 169 |
| TGCT | 0.8216 | 0.9426 | NA | 0.3559 | 0.6739 | NA | 139 | 137 | 2 |
| GBM | 0.8284 | 0.9426 | NA | 0.4795 | 0.7236 | NA | 602 | 528 | 74 |
| BRCA | 0.8827 | 0.971 | NA | 0.3198 | 0.6739 | NA | 1236 | 1136 | 100 |
| LUAD | 0.9689 | 0.9994 | NA | 0.251 | 1 | NA | 641 | 432 | 209 |
| LAML | 0.9966 | 0.9994 | NA | NA | NA | NA | 200 | 194 | 6 |
| HNSC | 0.9994 | 0.9994 | NA | 0.2647 | 0.6739 | 3282 | 604 | 519 | 85 |

<sup>†</sup> Table Key: FDR, False Discovery Rate; MUT, Desmosome coding mutations; WT, Desmosome wild type; NA, data not available.

**Supplementary Table 3:** Clinical and assay information on the 18 primary melanoma tumors^†^.

| Sample ID | Total mutations | WES | ST | IHC | Desmosome status | No. mutated desm. genes | Age | Sex | Tumor site | Thickness (mm) | Primary stage | pTNM | Stage |
| --- | --- | --- | --- | --- | --- | --- | --- | --- | --- | --- | --- | --- | --- |
| MKD1 | 334 | Y | Y | N | MUT | 1 | 40 | M | Back | 1 | pT1b | pT2aN1M0 | IIIA |
| MKD2 | 900 | Y | Y | N | MUT | 1 | 61 | M | Back | 1.1 | pT2a | pT2aN0M0 | IB |
| MKD3 | 368 | Y | Y | N | WT | 0 | 75 | M | Left arm | 0.9 | pT1b | pT1bNxMx | IA |
| MKD4 | 505 | Y | Y | N | MUT | 1 | 60 | M | Back | 2.1 | pT3b | pT3bNxMx | IIB |
| MKD5 | 1369 | Y | Y | N | MUT | 3 | 86 | F | Shoulder | 0.7 | pT1a | pT1aNxMx | IA |
| MKD6 | 304 | Y | Y | N | WT | 0 | 56 | M | Back | 0.5 | pT1a | pT1aNxMx | IA |
| MKD7 | 619 | Y | Y | N | WT | 0 | 54 | F | Right arm | 0.9 | pT1b | pT1aNxMx | IA |
| MKD8 | 420 | Y | Y | N | WT | 0 | 84 | F | Chest | 3.4 | pT3b | pT3bNxMx | IIB |
| MKD9 | 774 | Y | N | Y | MUT | 1 | 69 | F | Back | 14 | pT4a | pT4aNxMx | IIB |
| MKD10 | 1309 | Y | N | Y | WT | 0 | 48 | F | Shoulder | 1.9 | pT2a | pT2aNxMx | IB |
| MKD11 | 689 | Y | N | Y | WT | 0 | 66 | M | Scalp | 1.6 | pT2a | pT2aNxMx | IB |
| MKD12 | 1068 | Y | N | Y | MUT | 2 | 59 | F | Back | 1 | pT1b | pT1bNxMx | IB |
| MKD13 | 1233 | Y | N | Y | MUT | 3 | 57 | M | Abdomen | 1.4 | pT2a | pT2aNxMx | IB |
| MKD14 | 1368 | Y | N | Y | MUT | 3 | 50 | M | Forearm | 1.6 | pT2a | pT2aNxMx | IB |
| MKD15 | 1189 | Y | N | Y | WT | 0 | 79 | M | Back | 1.35 | pT2a | pT2aNxMx | IB |
| MKD16 | 1191 | Y | N | Y | MUT | 3 | 55 | F | Ear | 1 | pT1b | pT1bNxMx | IB |
| MKD17 | 1111 | Y | N | Y | WT | 0 | 74 | F | Arm | 6.3 | pT4b | pT4bNxMx | IIC |
| MKD18 | 1168 | Y | N | Y | WT | 0 | 75 | M | Forearm | 3.9 | pT3a | pT3aNxMx | IIA |

**† Table Key:** WES, Whole-Exome Sequencing; ST: Spatial Transcriptomics; IHC, Immunohistochemistry (Y = yes, N = no); pTNM, Pathological tumor-node-metastasis staging.

**Supplementary Table 4:** Genes used for cell proliferation signature. Related to Fig. 3d and Supplementary Fig. 7.

| Proliferation signature |  |  |  |  |  |
| --- | --- | --- | --- | --- | --- |
| ACP5 | CEACAM1 | HPS4 | NR4A3 | RRAGD | WDR91 |
| ADCY2 | DAPK1 | INPP4B | OCA2 | SEMA6A | ZFYVE16 |
| APOE | DCT | IRF4 | PHACTR1 | SIRPA |  |
| ASAH1 | FAM174B | IVNS1ABP | PIR | SLC45A2 |  |
| BIRC7 | GALNT3 | KAZ | PLXNC1 | ST3GAL6 |  |
| C21orf91 | GNPTAB | MBP | PMEL | STX7 |  |
| CAPN3 | GPM6B | MICAL1 | RAB27A | TNFRSF14 |  |
| CDH1 | GPR143 | MITF | RAB38 | TRPM1 |  |
| CDK2 | GPRC5B | MLANA | RGS20 | TYR |  |
| CDK5R1 | GYG2 | MYO1D | RHOQ | TYRP1 |  |

**Supplementary Table 5:**
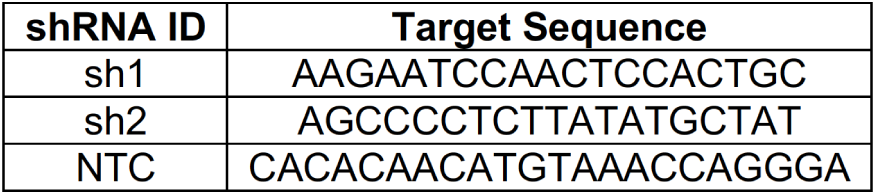
*DSP* shRNA sequences. Related to Figs. 6 and 7.

**Supplementary Fig. 1:**
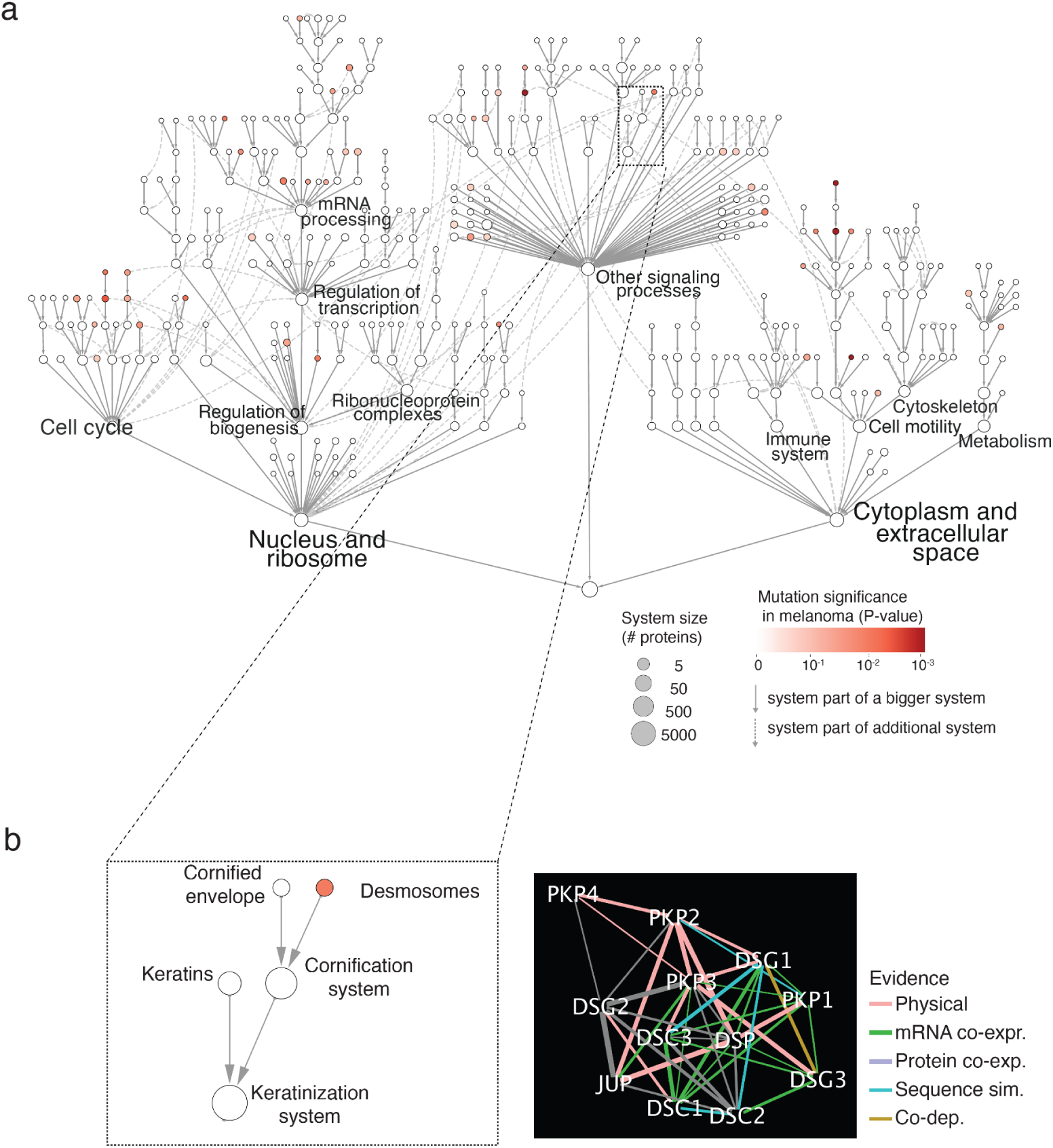
Interactions defining desmosome system and context in NeST hierarchy. (a) Nested Systems in Tumors (NeST) map showing all mutated systems identified across TCGA cohorts, published previously^17^. Nodes represent protein systems, identified in the previous study from dense communities of interacting proteins. Node size indicates the size of the system in number of proteins. Red intensity scale indicates significance of system mutation frequency in the skin cutaneous melanoma (SKCM) cohort. Gray arrows represent hierarchical containment (i.e. system 1 → system 2 denotes that system 1 is contained by system 2). For systems contained by multiple others (pleiotropy), each additional containment relation is connected by a dashed arrow. (b) Zoom detail of NeST hierarchy relevant to the desmosome system and its larger containing compartments (left). Adjacent is the specific protein interaction network that defines the desmosome system (right). Colors denote interaction types: Red: physical protein-protein interactions; green: correlation in mRNA expression; purple: correlation in protein abundance; cyan: protein sequence similarity; brown: co-dependency interactions, connecting proteins for which loss-of-function gene knockouts result in similar patterns of growth dependency across cell lines. See original publication^17^ for more details.

**Supplementary Fig. 2:**
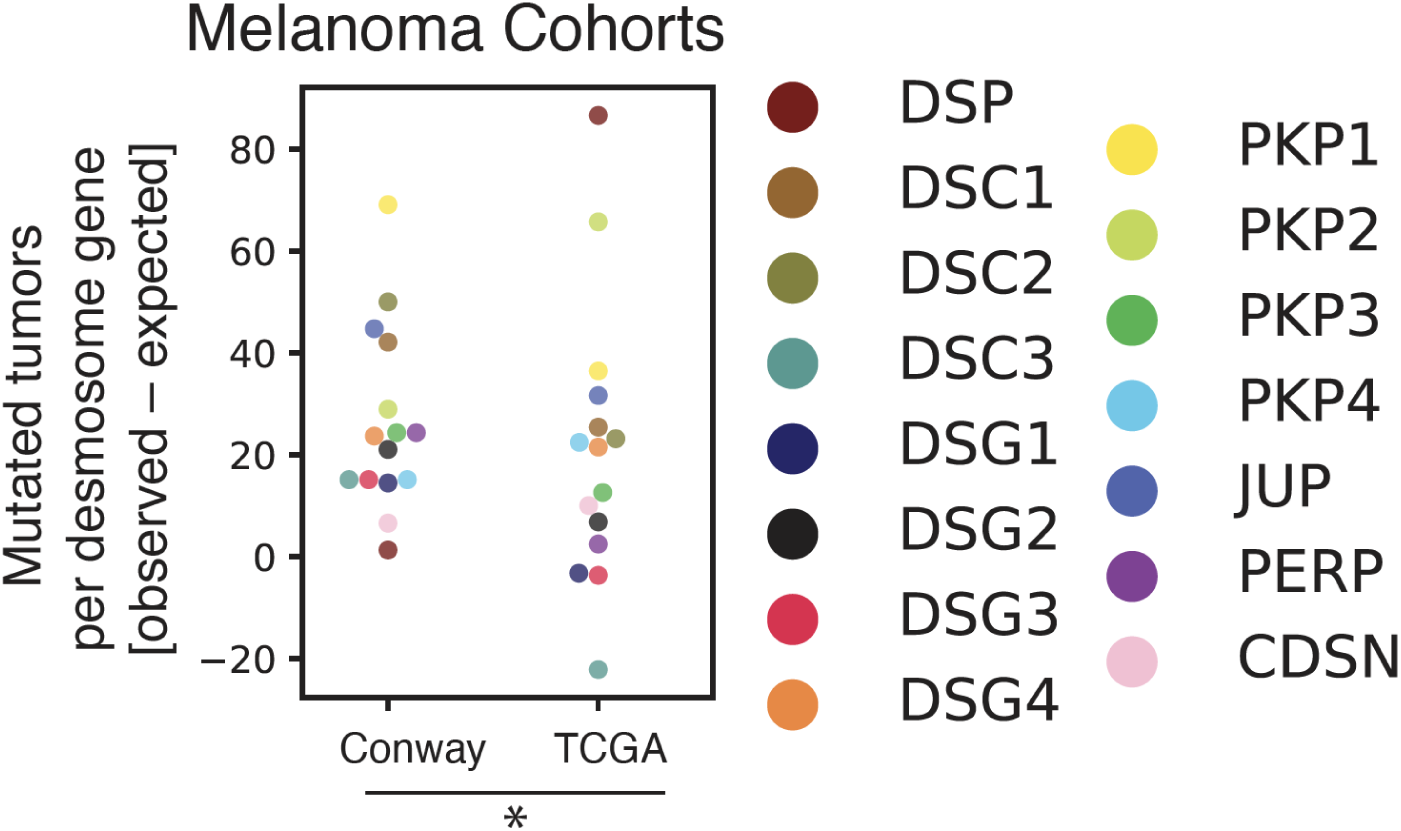
Desmosome mutation frequencies in multiple melanoma cohorts. Related to Fig. 1. Counts of mutated tumors per each desmosome gene (colored points) in the largest published melanoma cohort (Conway^20^, left) are compared to the SKCM-TCGA cohort (right). Horizontal black lines indicate median value across desmosomal genes. The set of desmosome genes has an above-expected number of mutations in both cohorts, with significance determined using bootstrap analysis (*, P<10^−10^). Desmosome subunits with consistently high mutation frequencies in both cohorts include PKP1, PKP2, DSC1, DSC2 and JUP. DSP has high mutation frequency in TCGA but not Conway. DSG1, DSG3 and DSC3 exhibit a low mutation frequency in both cohorts. Loss of these desmosome cadherins may not provide a selective advantage in melanoma due to functional redundancy with alternate cadherin genes.

**Supplementary Fig. 3:**
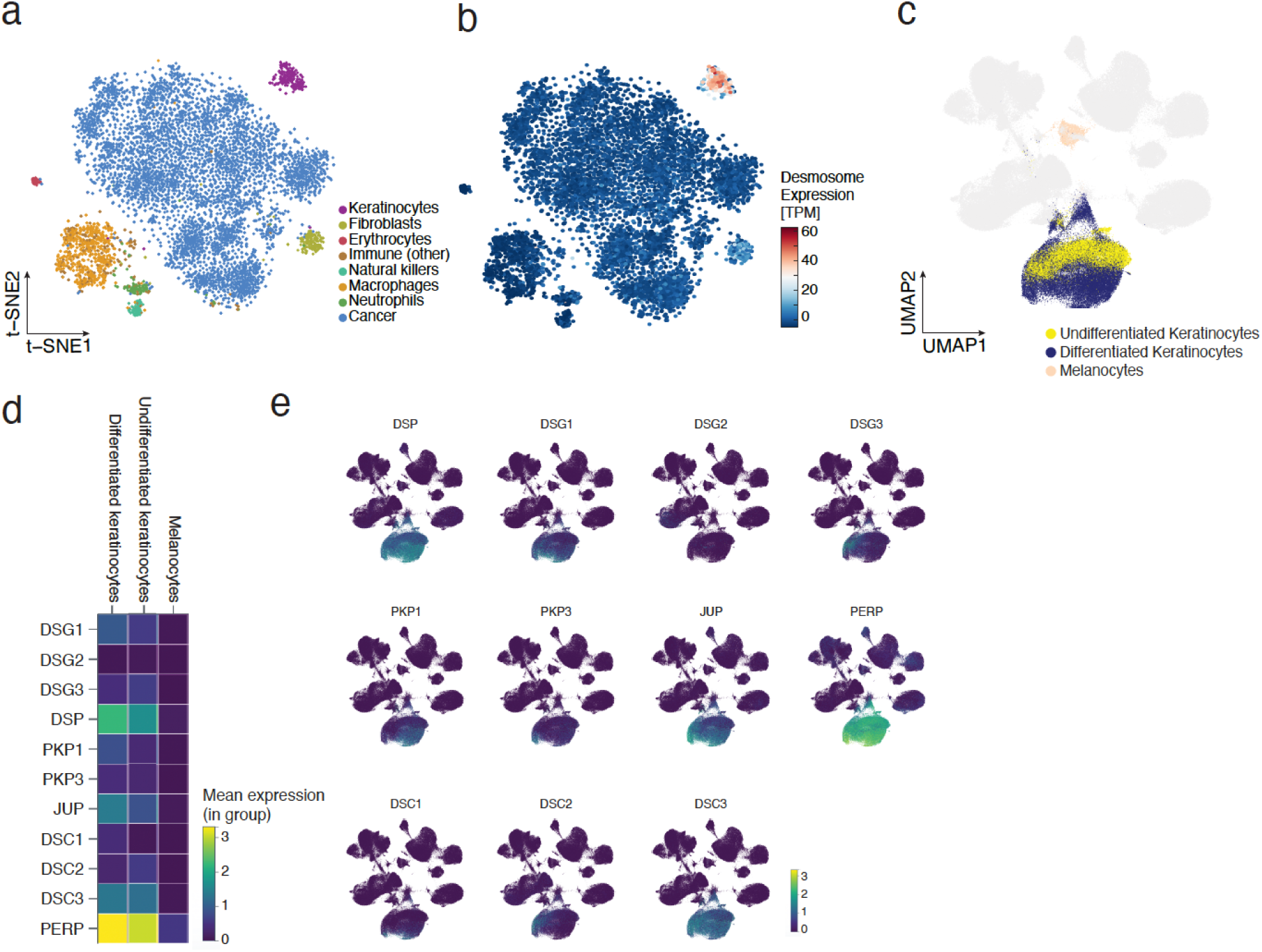
Single-cell RNA-seq analysis of desmosome gene expression in zebrafish melanoma samples and human skin. (a) t-distributed stochastic neighbor embedding (tSNE) analysis of approximately 7500 individual cells from zebrafish melanoma cells. Reanalysis of data published previously^36^. Color indicates inferred cell type. (b) Same tSNE plot as panel (a), colored by the expression of desmosome genes. (c) Uniform Manifold Approximation and Projection (UMAP) analysis of 239,045 individual cells from healthy human skin, with colors representing inferred cell types of interest (undifferentiated keratinocytes, differentiated keratinocytes, melanocytes). Uncolored (gray) dots correspond to other cell types found in skin including fibroblasts, vascular cells, and lymphocytes.. (d) Heatmap showing the average expression levels of desmosome genes in melanocytes as well as differentiated and undifferentiated keratinocytes. (e) Grid of UMAPs from (c), each colored by expression of a different desmosome gene.

**Supplementary Fig. 4:**
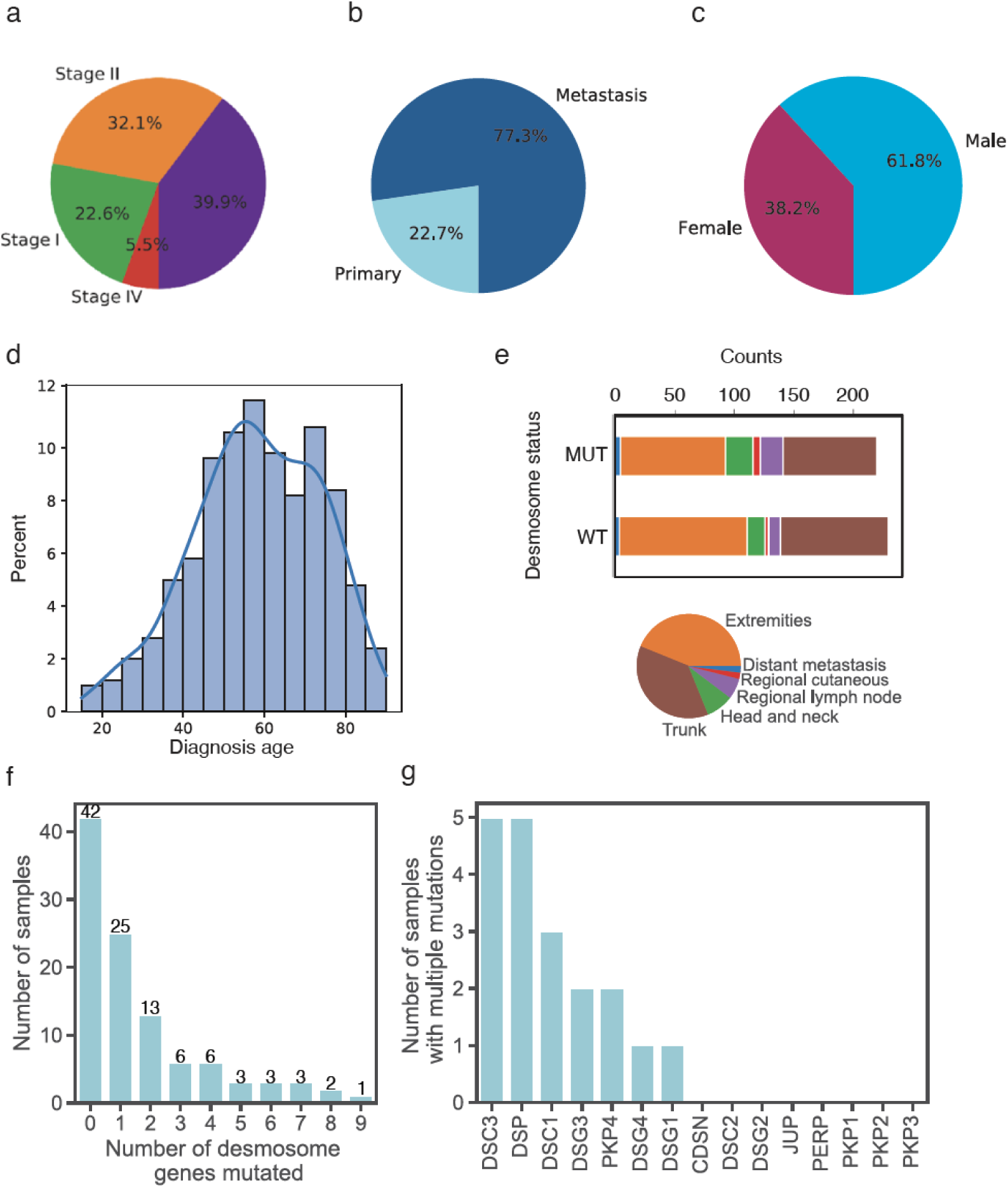
TCGA-SKCM cohort summary information. Shown for N = 472 samples in the TCGA-SKCM cohort. (a) Pie chart describing clinical staging distribution. (b) Pie chart describing sample type distribution. (c) Pie chart describing sex distribution. (d) Histogram describing diagnosis age distribution. (e) Lower: Pie chart describing tumor site distribution. Upper: Tumor site distribution is divided in separate bar plots shown for high desmosome mutation burden (MUT) versus low desmosome mutation burden (WT). MUT and WT are defined by whether the desmosome mutation count is above or below the median value across all patients, respectively. (f) Bar plot depicting the distribution of the number of mutated desmosome genes across primary melanoma tumor samples in the SKCM-TCGA cohort. (g) Bar plot depicting the distribution of primary tumors with multiple coding mutations across desmosome genes in the SKCM-TCGA cohort.

**Supplementary Fig. 5:**
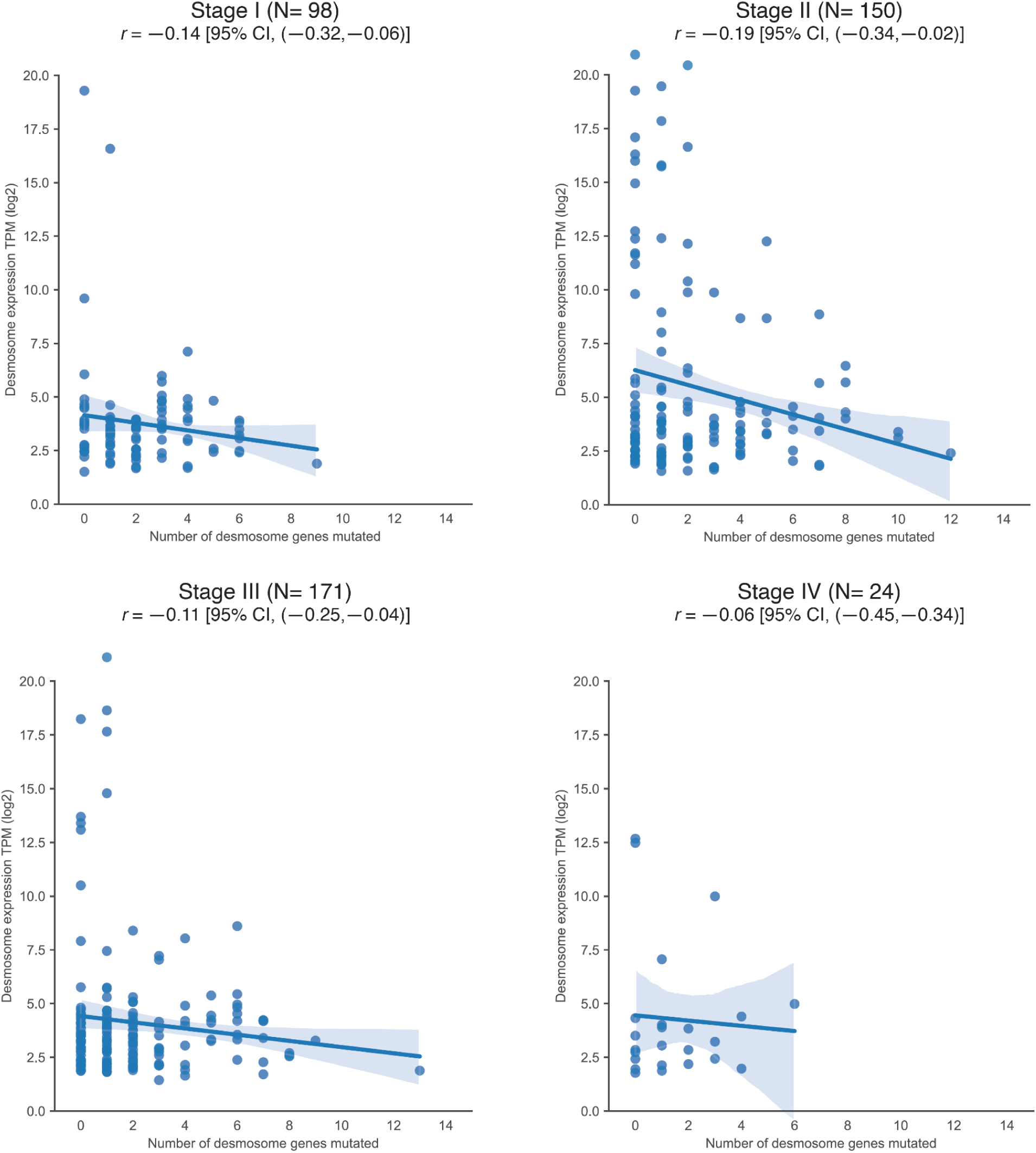
Relationship of desmosome expression to number of desmosome mutations across tumor stages. Related to Fig. 3c. Average desmosome gene expression (TPM, log2) is shown as a function of the number of desmosome genes with coding mutations. Plots are shown separately for stage I (top left), stage II (top right), stage III (bottom left) and stage IV (bottom right) melanoma tumor populations. Regression lines for each of these populations are shown. Shaded areas capture 95% confidence intervals. Title for each panel shows Pearson correlation coefficients with corresponding confidence intervals and total numbers of samples (N).

**Supplementary Fig. 6:**
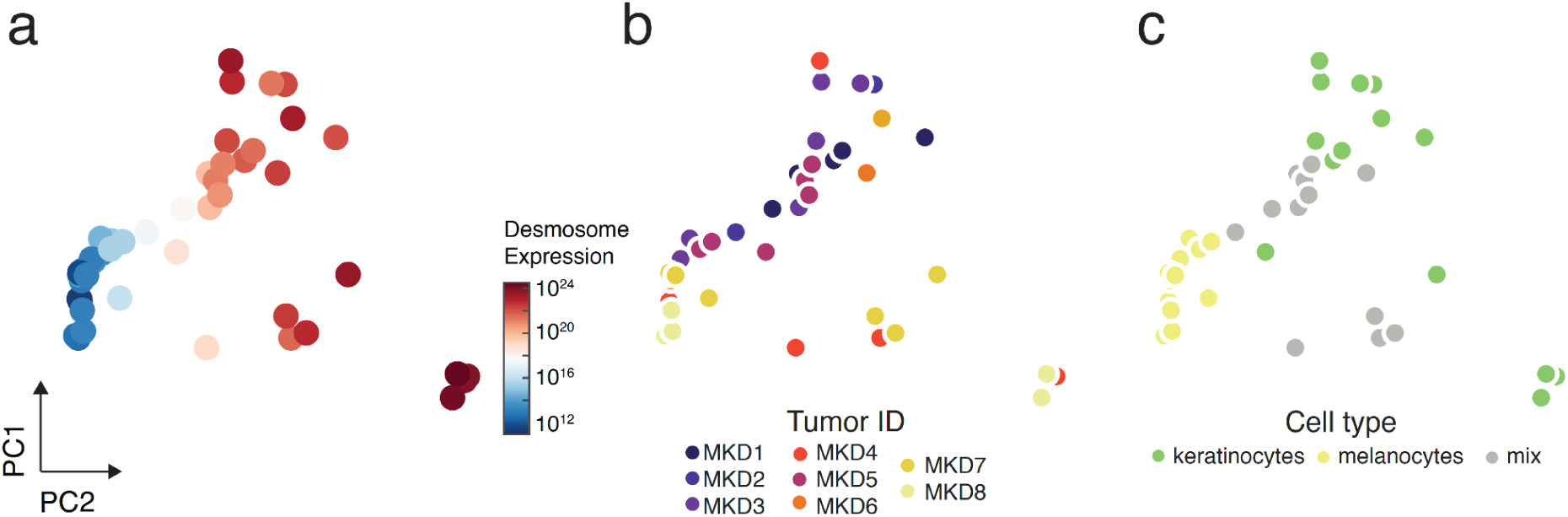
Principal Component Analysis (PCA) of spatial transcriptomic data. Related to Fig. 4. (a) Profiles of mRNA expression for each ROI (covering 18,695 genes total) are reduced to the first two principal components (PC1 versus PC2). A total of 48 ROIs are drawn from 8 tumor biopsies. Color indicates average expression of desmosome genes. (b) Same PCA as (a), color indicates tumor ID. (c) Same PCA as (a), color indicates cell type identified by antibody staining.

**Supplementary Fig. 7:**
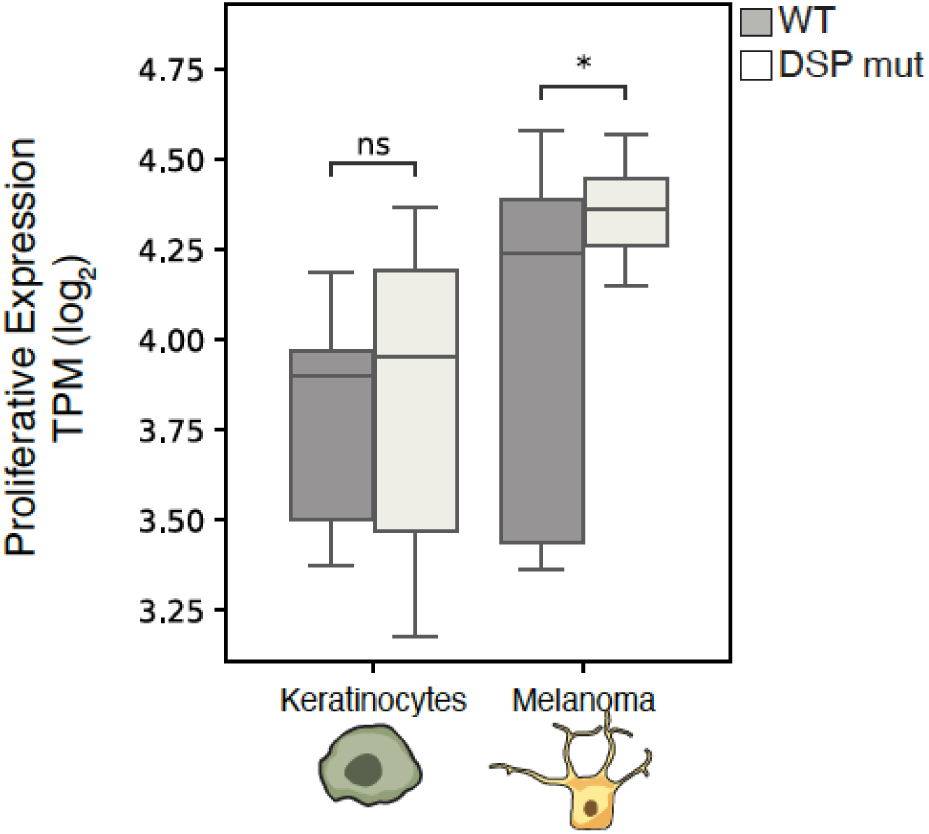
DSP mutations show increase in the proliferative gene expression program. Related to Fig. 4. Degree of proliferative gene expression (as defined previously^70^, see also Supplementary Table 4) in desmosome WT (dark gray, N = 32) or desmosome mutant (light gray, N = 16) regions of interest (ROI) from primary melanoma tumors in the spatial transcriptomics cohort. Significance determined by two sample t-test (*, P<0.05). The boxes contain the 25th to 75th percentile, the middle line denotes the 50th percentile, and the whiskers mark the 5th and 95th percentiles.

**Supplementary Fig. 8:**
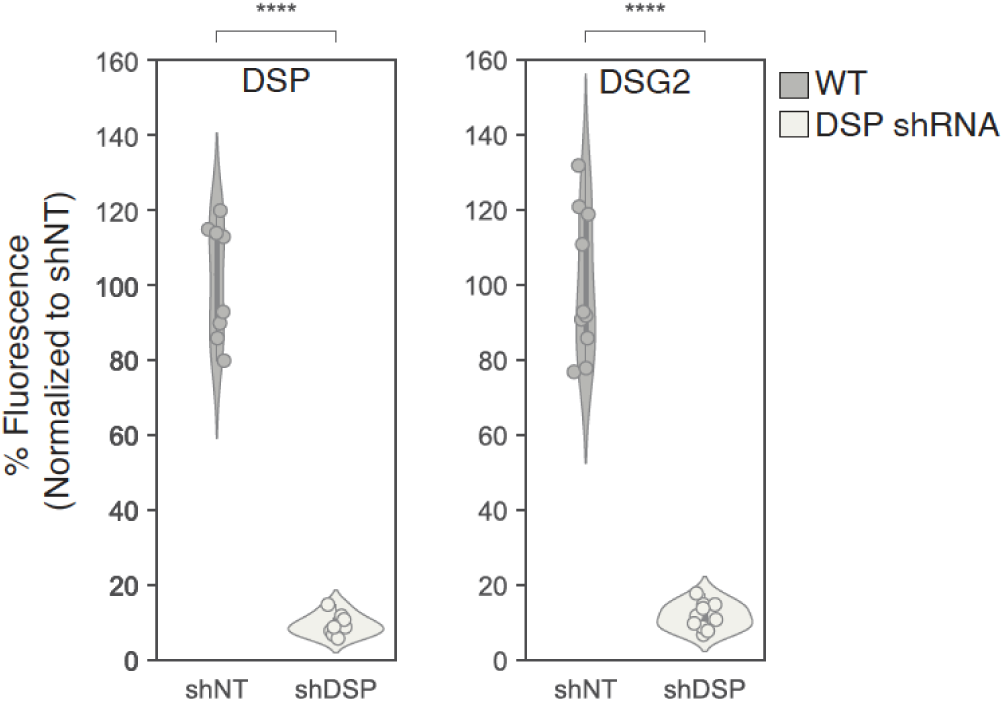
Immunofluorescence assay to measure desmosome protein abundance. Related to Fig. 6. *DSP* (left) and *DSG2* (right) relative protein expression levels in keratinocytes stably expressing NTC (dark gray, N = 9 replicates) or *DSP* shRNA (light gray, N = 9 replicates). Significance determined based on two sample t-test (****, P<0.0001).

**Supplementary Fig. 9:**
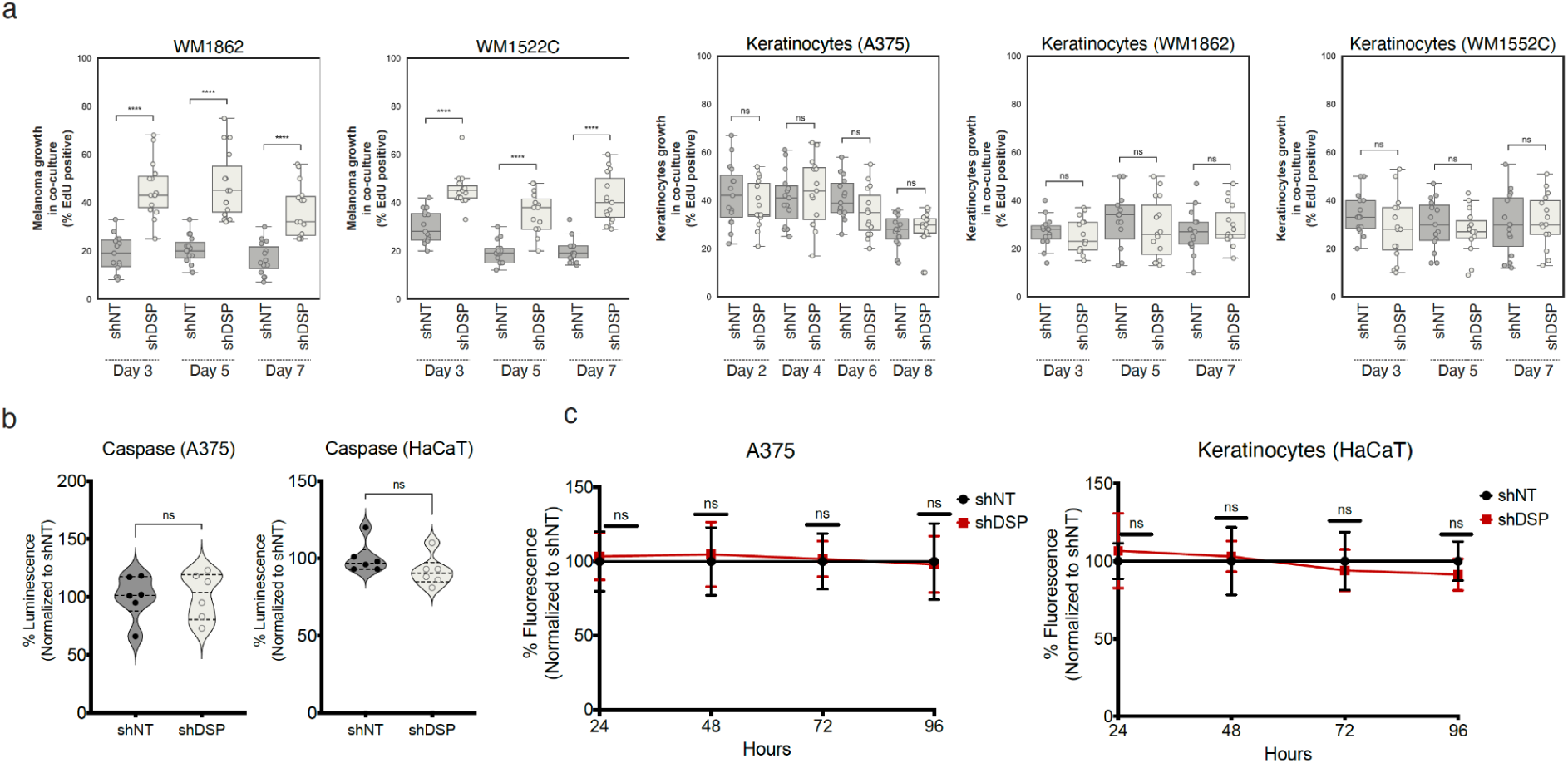
Effects of *DSP* disruption on proliferation using primary melanoma lines. Related to Fig. 7. (a) EdU proliferation measurements involving additional melanoma cell lines and keratinocytes, shown for co-cultures in which keratinocytes stably express NTC shRNA (dark gray) versus *DSP* shRNA (light gray) for 2, 4, 6 or 8 days, or 3, 5 or 7 days (indicated on x-axis). Each box summarizes 5 distinct fields per well ⨉ 3 biological replicates for N = 15 replicate co-culture measurements. Significance based on two sample t-test (**** P<0.0001, ns - not significant). The boxes contain the 25th to 75th percentile, the middle line denotes the 50th percentile, and the whiskers mark the 5th and 95th percentiles. The cell type (melanoma cell line or HaCaT keratinocyte) measured for proliferation is shown above each plot. For keratinocytes, the co-cultured melanoma cell line is shown in parentheses. (b) Caspase Glo assay for measurement of apoptosis in melanoma cells (A375, left) or keratinocytes (HaCaT, right). Significance based on two sample t-test (ns - not significant). (c) Cyquant assay of melanoma and keratinocyte cell proliferation measurements at 24, 48, 72 and 96 hours.

**Supplementary Fig. 10:**
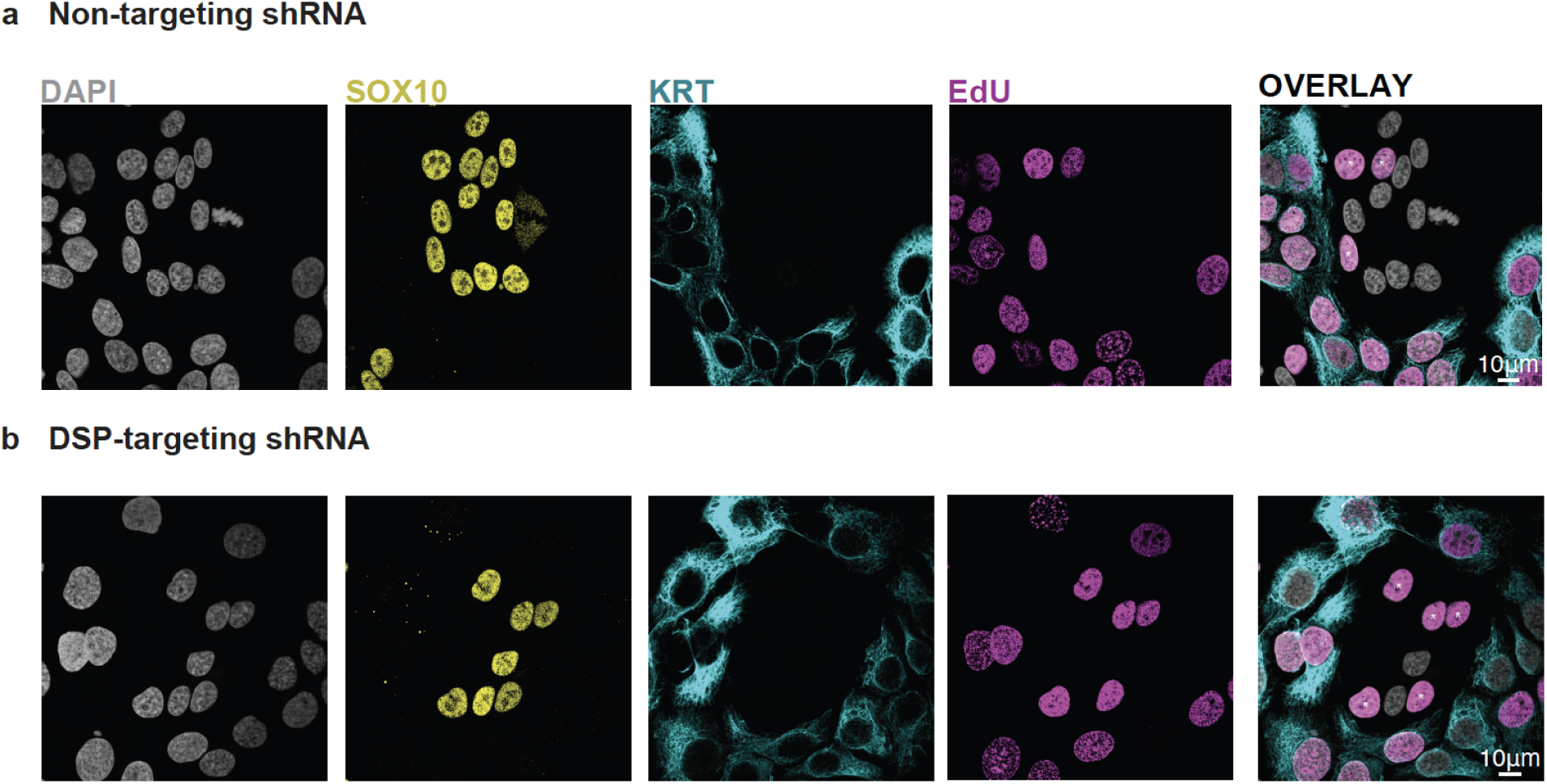
Effects of *DSP* gene disruption in melanoma/keratinocyte co-cultures. Related to Fig. 7. (a) Representative images from the melanoma/keratinocyte co-cultures with keratinocytes expressing stable NTC shRNA, depicting staining for SOX10 (yellow, melanoma marker), KRT14 (cyan, keratinocyte marker) and EdU (purple, proliferation marker). Grey color represents all cell nuclei (DAPI). First four panels display each staining individually, while the final right-most panel presents a merged image overlay. The white asterisk indicates proliferating melanoma cells (SOX10^+^,EDU^+^). (b) Same as (a) but from the melanoma/keratinocyte co-cultures with keratinocytes expressing stable DSP shRNA.

**Supplementary Fig. 11:**
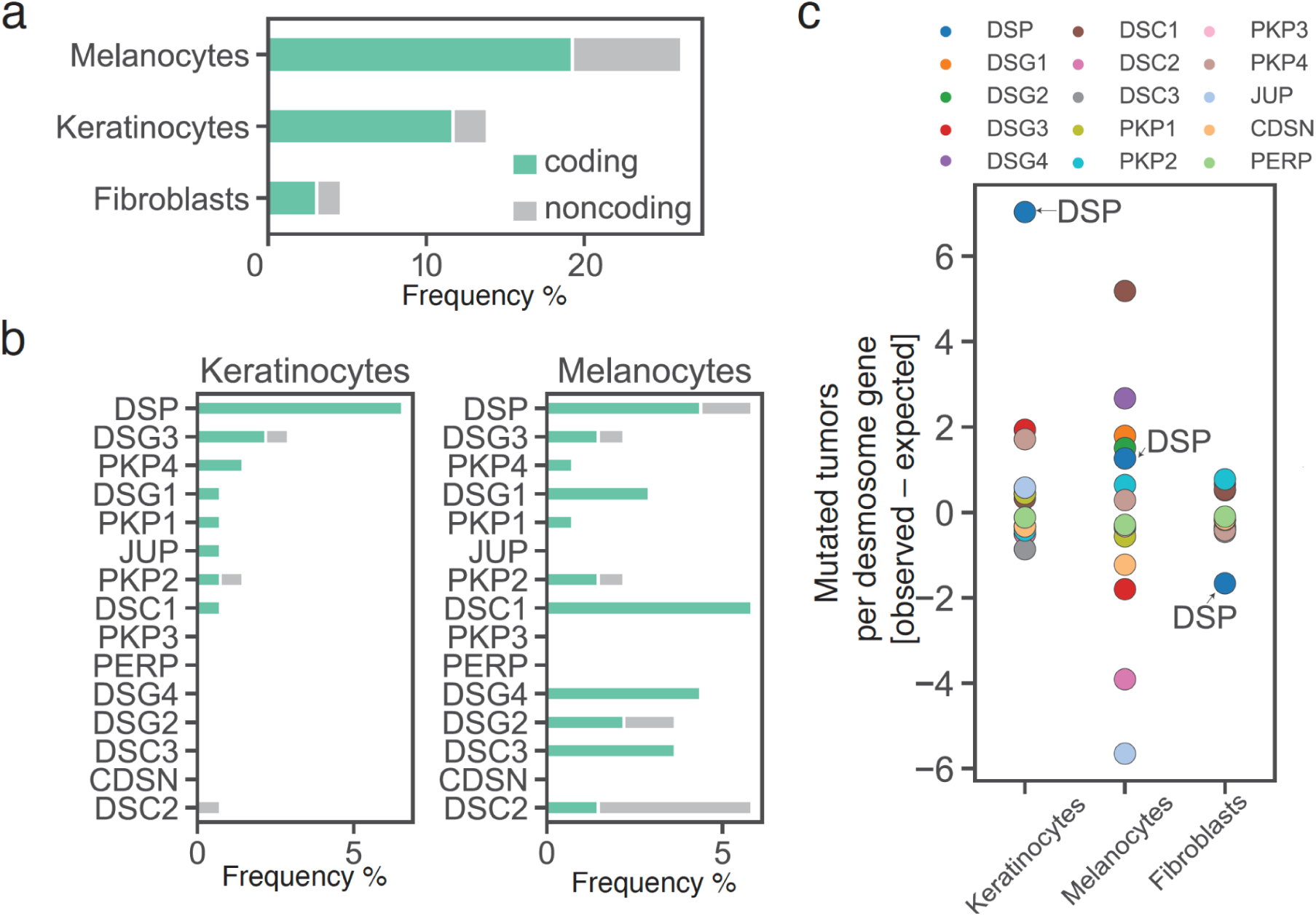
Desmosome mutational patterns in a single-cell DNA sequencing cohort. Whole-exome DNA sequences of clonally-expanded normal skin cells were collected in a recent preprint^54^, totalling 137 keratinocytes, 130 melanocytes and 23 fibroblasts collected from 15 donors. (a) Frequency of desmosome mutations for each of the three skin cell types sequenced in the cohort, aggregated across 15 desmosome genes. The stacked bars indicate the proportion of coding (green) versus non-coding (gray) mutations. (b) Same as (a) but for each desmosome gene, shown separately for keratinocytes (left) and melanocytes (right). (c) Excess number of cells with a mutated desmosome gene (colored points) above the number expected by chance (Online Methods). Similar to Fig. 1c. The genome-wide background mutation rate is substantially higher for melanocytes than keratinocytes (approximately 5 versus 1 mutation/Mb, as noted by the study authors^54^). Correcting for this expected background, the highest burden of desmosomal mutations is seen in keratinocytes, particularly for the gene DSP.

## Notes

### Summary of Updates

A major new body of work providing immunofluorescence results for 10 of our original melanoma tumor samples (new Fig. 5). These data were generated to robustly validate our earlier finding that desmosome mutations are associated with reduced desmosome protein expression specifically in keratinocytes. The new results also confirm that this effect is localized to the cell membrane, as expected for desmosomes, and does not extend to other compartments such as the cytoplasm. We have performed additional single-cell RNA sequencing analysis of over 200,000 cells from the Human Cell Atlas skin dataset, to confirm our earlier findings that desmosome expression is enriched in keratinocytes compared to melanocytes. We have included new Discussion paragraphs to discuss these and various other points suggested by these same reviewers. We have analyzed a late-breaking dataset by the Shain lab, which makes available genomic DNA sequences for clonally amplified keratinocytes and melanocytes isolated from normal human skin samples. This analysis does support the presence of desmosome mutations in keratinocytes.

